# Identification of a novel *chalcone reductase* gene for isoliquiritigenin biosynthesis in dahlia (*Dahlia variabilis*)

**DOI:** 10.1101/2022.03.28.486017

**Authors:** Sho Ohno, Haruka Yamada, Kei Maruyama, Ayumi Deguchi, Yasunari Kato, Mizuki Yokota, Fumi Tatsuzawa, Munetaka Hosokawa, Motoaki Doi

## Abstract

Butein is one of flavonoids conferring bright yellow flower color and is a precursor of aurone in some species. Butein is synthesized by two steps, 3-malonyl CoA and 4-coumaloyl CoA are converted to isoliquiritigenin in the first step, and then isoliquiritigenin is converted to butein in the second step. In the first step, chalcone synthase (CHS) and chalcone reductase (CHR) catalyze this reaction, however, CHR has been reported for the isoflavone biosynthesis pathway in legumes, and CHR for butein biosynthesis has not yet been isolated. In this study, we report CHR that is evolutionally different gene from legume species is involved in isoliquiritigenin biosynthesis in dahlia. To isolate CHR gene, we conducted comparative RNA-seq analysis between ‘Shukuhai’ and its butein-loss lateral mutant ‘Rinka’. We found *DvCHR* showed significant difference in expression levels that encodes an aldo-keto reductase (AKR) 13 family protein, which was phylogenetically different from legume CHRs belonging to AKR4A family. Gene expression levels and genotype of *DvCHR* were correlated with butein accumulation among various dahlia cultivars. Though single over expression of *DvCHR* was not able to accumulate isoliquiritigenin in tobacco, co-overexpression of *DvCHR* with a chalcone glucosyltransferase *Am4′CGT* and a MYB transcription factor *CaMYBA* successfully induced isoliquiritigenin accumulation. In addition, *DvCHR* homologous gene expression was detected from butein or aurone accumulating Coreopsideae species but not from non-butein or non-aurone accumulating *Asteraceae* species. These results indicated *DvCHR* functions as chalcone reductase for butein biosynthesis in dahlia, and isoliquiritigenin biosynthesis in Coreopsideae species has been developed independently from legume species.

## Introduction

Flower color is one of the most important traits in floriculture industry. Among various flower colors, yellow coloration is mainly derived from accumulation of carotenoids or betaxanthins, thus, species that are not able to biosynthesize and accumulate these pigments in flowers rarely have bright yellow flower. There are a few flavonoids known to exhibit yellow coloration. One of yellow flavonoids is butein (2′,3,4,4′-tetrahydroxychalcone), which exhibits bright yellow color in flowers and detected in limited species such as in dahlia (Price, 1939), *Cosmos sulphureus* (Geissman, 1942) and *Coreopsis grandiflora* (Geissman and Heaton, 1943). Butein is presumed to be synthesized from common substrate of flavonoids, malonyl-CoA and 4-coumaroyl-CoA, which are converted to 2′,4,4′-trihydroxychalcone (isoliquiritigenin or 6′-deoxychalcone) by chalcone synthase (CHS) and chalcone reductase (CHR), and then hydroxylated by chalcone 3-hydroxylase (CH3H) (Fig. 1). A gene encoding CH3H has been elucidated in *C. sulphureus* (Schlangen et al, 2010a), and an allelic variant of flavonoid 3′-hydroxylase (F3′H) in dahlia also had a function as chalcone 3-hydroxylase activity (Schlangen et al, 2010b). Another yellow flavonoid is aurone, which is found in snapdragon (*Antirrhinum majus*) and some Asteraceae species (Miosic et al, 2013). Aurone is synthesized by two key enzymes [aureusidin synthase (AS) and chalcone 4′-*O*-glucosyltransferase (Am4′CGT)] in snapdragon, whereas aurone is synthesized from butein by a polyphenol oxidase in *Coreopsis* (Kaintz et al, 2014; Molitor et al, 2015: Fig. 1). In this butein or aurone biosynthetic pathway, only CHR has not been identified yet.

**Fig. 1.**
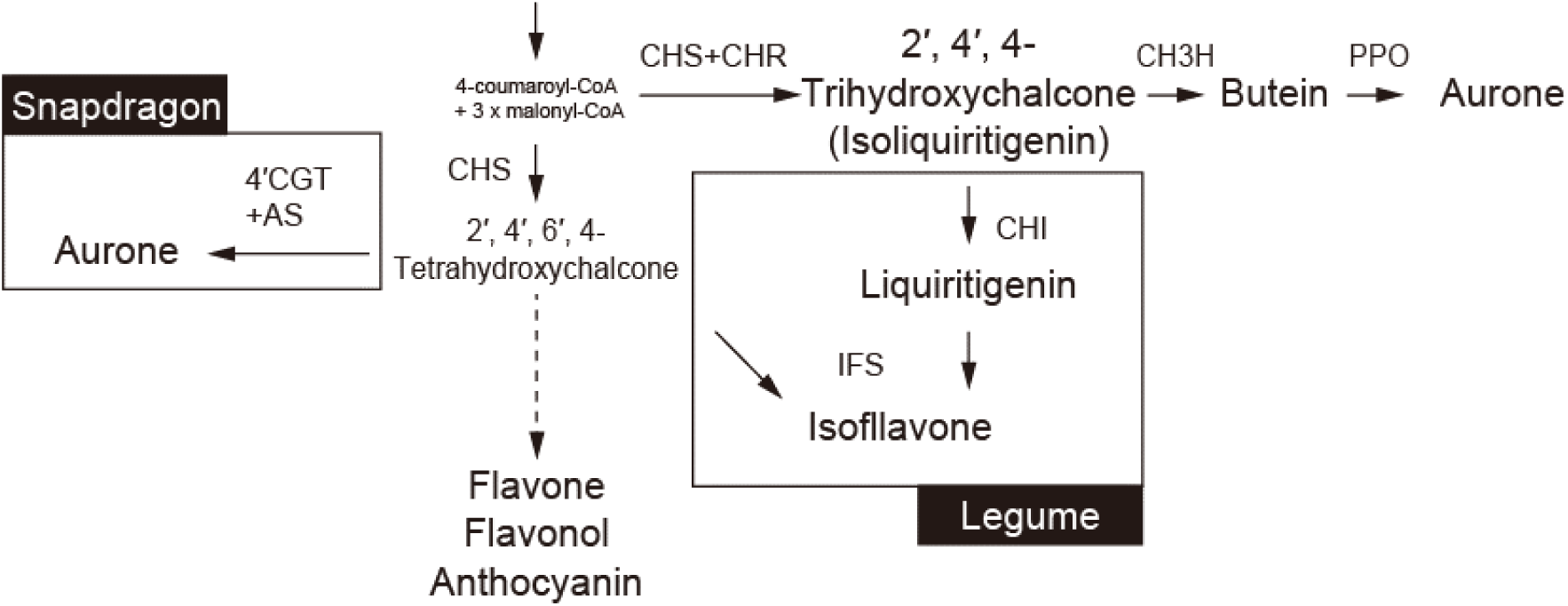
Simplified flavonoid biosynthesis pathway. Abbreviations: AS, aureusidin synthase; CH3H, chalcone 3-hydroxylase; CHI, chalcone isomerase; CHR, chalcone reductase; CHS, chalcone synthase; IFS, isoflavone synthase; PPO, polyphenol oxidase; 4′CGT, chalcone 4′-*O*-glucosyltransferase.

Dahlias (*Dahlia variabilis*) are popular ornamental plants which belong to Asteraceae and are autoallooctoploids with chromosome number 2n = 8x = 64 (Gatt et al, 1998). Dahlia flowers exhibit huge variations in flower color by flavonoid pigments, and many genes associated with flavonoid biosynthesis in ray florets were analyzed (Suzuki et al, 2002; Ohno et al, 2011a; 2011b; 2013b; 2018a; 2021; Deguchi et al, 2013). Although dahlias are able to biosynthesize only flavonoids and do not accumulate carotenoid or betaxanthins in ray florets, there are a lot of bright yellow flower cultivars. The pigment contributes to yellow flower color in dahlia is a butein (Price, 1939), and the final products were determined as butein 4’-malonylsophoroside and butein 4’-malonylglucoside (Harborne et al, 1990). Butein derivatives were also detected in leaves (Ohno et al, 2018b). In dahlia, butein always co-accumulates with isoliquiritigenin, suggesting CHR is the determinant factor for butein biosynthesis (Ohno et al, 2013a).

CHR is characterized only in legumes for isoflavone synthetic pathway. CHR activity was at first found in *Glycyrrhiza echinata* (Ayabe et al, 1998) and in soybean (*Glycine max*) (Welle and Grisebach, 1988), and then cDNA sequence of *GmCHR1* was determined in soybean (Welle et al, 1991). *GmCHR* and all other legume CHRs associated with isoliquiritigenin biosynthesis in isoflavone biosynthesis pathway belongs to aldo-keto reductase (AKR) 4A sub-family (Jez et al, 1997; Bomati et al, 2005). However, in dahlia, no orthologous AKR 4A sub-family gene was found by RNA-seq analysis in a preliminary experiment, suggesting a gene encoding CHR in dahlia is completely different from *CHR* in legumes. Thus, in this study, to explore a candidate gene encoding CHR for butein biosynthesis in dahlia, comparative transcriptome analysis was conducted using lateral mutant cultivars that had difference in butein biosynthesis. We identified one contig named *DvCHR* which belongs to AKR 13 sub-family, and confirmed the function of *DvCHR* by transgenic approach.

## Results

### ‘Rinka (RK)’ loses isoliquiritigenin and butein biosynthesis in ray florets

In this study, we used a red-white bicolor dahlia cultivar ‘Shukuhai (SH)’ and its lateral mutant cultivars, ‘Iwaibune (IB)’ and ‘Rinka (RK)’ (Fig 2A). ‘SH’ accumulates anthocyanins, flavones and chalcones (butein and isoliquiritigenin) in the red part of ray florets. ‘IB’, a lateral mutant from ‘SH’, has deeper red color than ‘SH’ and accumulates the same three pigments but higher amount of anthocyanins and flavones and lower amount of chalcones (Fig 2C). ‘RK’, a lateral mutant from ‘IB’, produces purple flowers and accumulates anthocyanins and flavones but no chalcones (Fig 2D). Along with mutations from ‘SH’ to ‘IB’ or ‘IB’ to ‘RK’, gradual reduction of chalcones were observed. Since butein is presumed to be biosynthesized from isoliquiritigenin, therefore, it was suggested that mutations had been occurred in the genes that are involved in isoliquiritigenin biosynthesis in these lateral mutant cultivars ‘IB’ and ‘RK’.

**Fig. 2.**
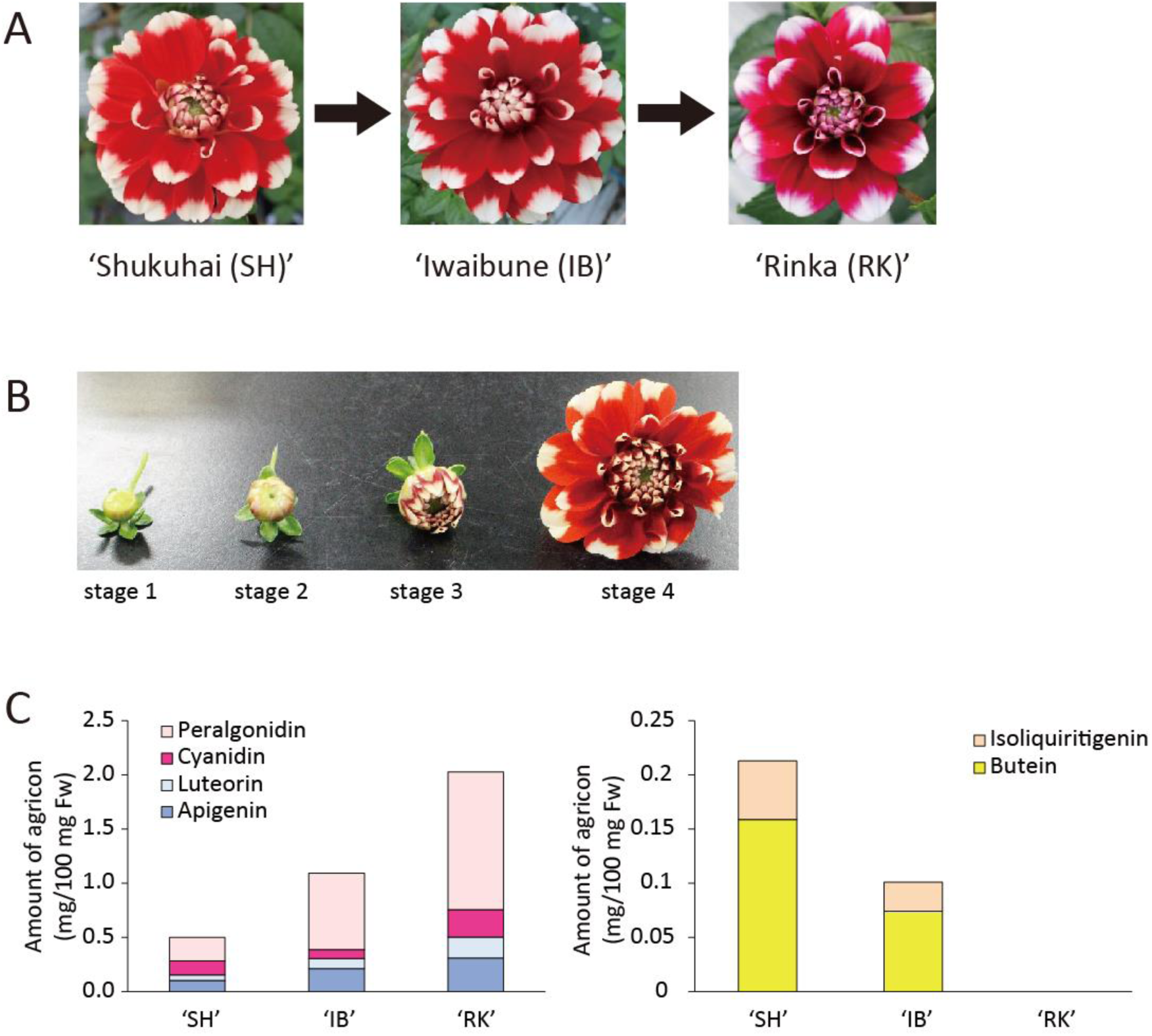
Pigment analysis among ‘Shukuhai’, ‘Iwaibune’ and ‘Rinka’. A, Dahlia cultivars used in this study. B, Ray floret developmental stages used in this study. C, Anthocyanin and flavone content at the stage 3 ray florets. and D, Butein and isoliquiritigenin content at the stage 3 ray florets.

### CHS, CH3H and AKR 4B sub-family genes are unlikely to the determinant factor for loss of isoliquiritigenin and butein biosynthesis in ‘Rinka (RK)’

At first, we examine *DvCHS2* and *DvCH3H* genes. Previous studies suggested *DvCHS2* plays a role in anthocyanin, flavone and butein biosynthesis in dahlia (Ohno et al, 2011a; 2018a). Since CH3H gene in dahlia had not been isolated, we isolated *DvCH3H* which is paralogous to *CsCH3H* gene which shares 90% homology and contains substrate recognition site (SRS1) and XSAGGXX domain (Fig. S1; Schlangen et al., 2010b). Schlangen et al. (2010a) reported that one of allelic variants of *F3′H* showed chalcone 3-hydroxylase activity, and they indicated the valine at position 425 was decisive for chalcone acceptance. DvCH3H also carries the valine at position 425 (Fig. S1). Next, we analyzed expression levels of *DvCHS2* and *DvCH3H*, but there was no difference between ‘SH’ and ‘RK’ (Fig. S2). We analyzed transcript sequence of *DvCHS2* and *DvCH3H*, two sequences were isolated for *DvCHS2,* and one sequence was isolated for *DvCH3H*, however no difference was found between ‘SH’ and ‘RK’. These results suggested that *DvCHS2* and *DvCH3H* is not the determinant factor of butein biosynthesis in these cultivars. Therefore, CHR would be the causal gene for loss of isoliquiritigenin and butein biosynthesis in ‘RK’.

CHR genes isolated so far are only from legumes for isoflavone biosynthetic pathway belong to AKR 4A sub-family (Fig. 3A). This family is consisted of only legume CHRs and does not include any other aldo-keto reductase genes. We tried to isolate AKR 4A sub-family genes expressing in the ray florets by RNA-seq, but we could not find any related genes. Instead, the phylogenetically closest AKR genes expressing in ray florets are four AKR 4B sub-family genes (*DvAKR1* - *DvAKR4*). We compared gene expression of these genes among ‘SH’, ‘IB’ and ‘RK’, however there was no significant difference (Fig. S2). From these results, it was considered that the gene involved in the isoliquiritigenin biosynthesis in dahlia is a novel type of CHR and is totally different from legume CHRs.

**Fig. 3.**
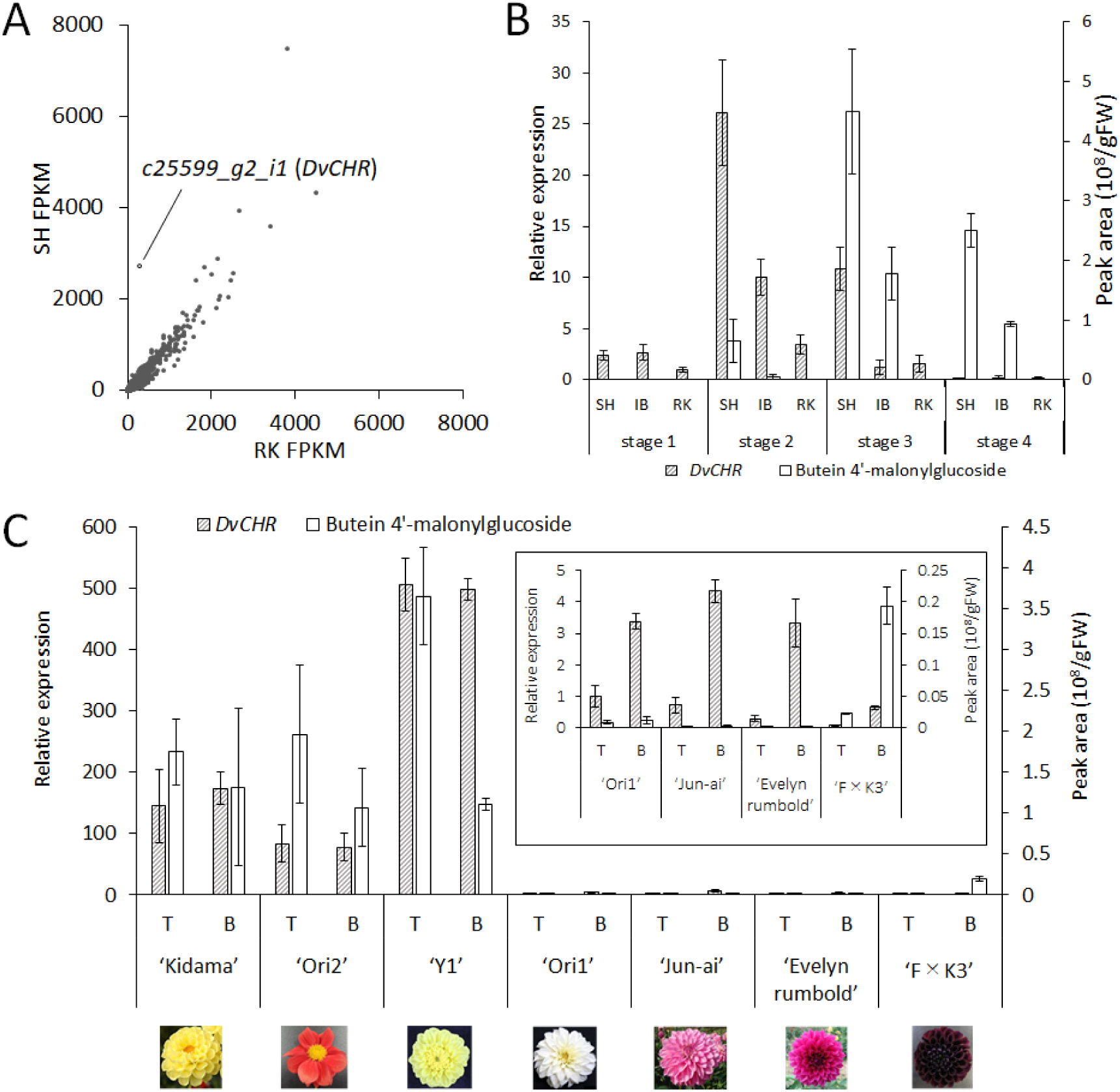
*DvCHR* expression analysis. A, Scatter plot of transcriptome between ‘SH’ and ‘RK’ Fragments Per Kilobase of exon per Million mapped reads (FPKM). B, Relative expression of *DvCHR* and peak area of butein 4’-malonylglucoside measured by HPLC in ray florets of ‘SH’, ‘IB’ and ‘RK’. C, Relative expression of *DvCHR* and peak area of butein 4’-malonylglucoside measured by HPLC in ray florets of various dahlia cultivars. T: top half, B: bottom half. For relative *DvCHR* expression, expression level of ‘Ori1’ top half was calculated as one. *DvActin* was used as an internal control. Bars represent standard error (n = 3).

### Comparative RNA-seq analysis indicates c25599_g2_i1 is a CHR candidate gene

To explore the CHR gene in dahlia, at first, we conducted comparative RNA-seq analysis between ‘SH’ and ‘RK’ ray florets at stage 2. Total 106,692 contigs were constructed and 74,045 of these contigs were translated into amino acid sequences by *de novo* assemble. Among top 1000 contigs in mean FPKM value of ‘SH’ and ‘RK’, only one contig (*c25599_g2_i1*) showed more than three-fold difference between ‘SH’ and ‘RK’. While FPKM value of other genes related to flavonoid biosynthesis including *DvCHS1*, *DvCHS2*, *DvCH3H* were less than twice higher or lower between ‘SH’ and ‘RK’ (Table S1), FPKM value of *c25599_g2_i1* was approximately 9.5 times higher in ‘SH’ than in ‘RK’ (Fig. 3A; Table S1). Thus, we named this *c25599_g2_i1* as *DvCHR*. Next, we analyzed correlation of *DvCHR* expression level and butein accumulation in ray florets at different developmental stages. Among ‘SH’, ‘IB’ and ‘RK’, *DvCHR* expression pattern was correlated to accumulation of butein 4′-malonylglucoside and relatively low expression level was detected in ‘RK’ (Fig. 3B). In addition, we analyzed other cultivars with various flower colors. We separated stage 2 ray florets into top half and bottom half, because most of cultivars accumulates small content of butein in the basal half part, but only yellow or red cultivars accumulate butein in the top half part. Butein accumulating cultivars (‘Kidama’, ‘Ori2’ and ‘Y1’) exhibited significantly elevated *DvCHR* expression compared with those of butein less-accumulating cultivars (‘Ori1’, ‘Jun-ai’, ‘Evelyn Rumbold’ and ‘F×K3’) correlated with butein 4′-malonylglucoside (Fig. 3C). Therefore, these results supported the hypothesis that *DvCHR* is a candidate gene for *CHR* in dahlia.

The full-length ORF of *DvCHR* CDS was 1065 bp encoding 354 amino acid residues. BLAST search (http://blast.ncbi.nlm.nih.gov/Blast.cgi) revealed that *DvCHR* belongs to AKR superfamily, but far from legume CHRs that belong to AKR 4A sub-family (Fig. 4A). *DvCHR* shares relatively high homology with AKR 13 family such as *Arabidopsis* AT1G60710 (69%), *Rauvolfia serpentine* perakine reductase (55%) and *Perilla frutescens* alcohol dehydrogenase (64%) (Fig. 3B). DvCHR contains conserved catalytic tetrad (Asp^57^, Tyr ^62^, Lys^88^, His^130^) and several typical AKR cofactor binding residues (Ser^160^, Gln^180^ and Tyr^208^) (Fig. 3B: Sun et al., 2012).

**Fig. 4.**
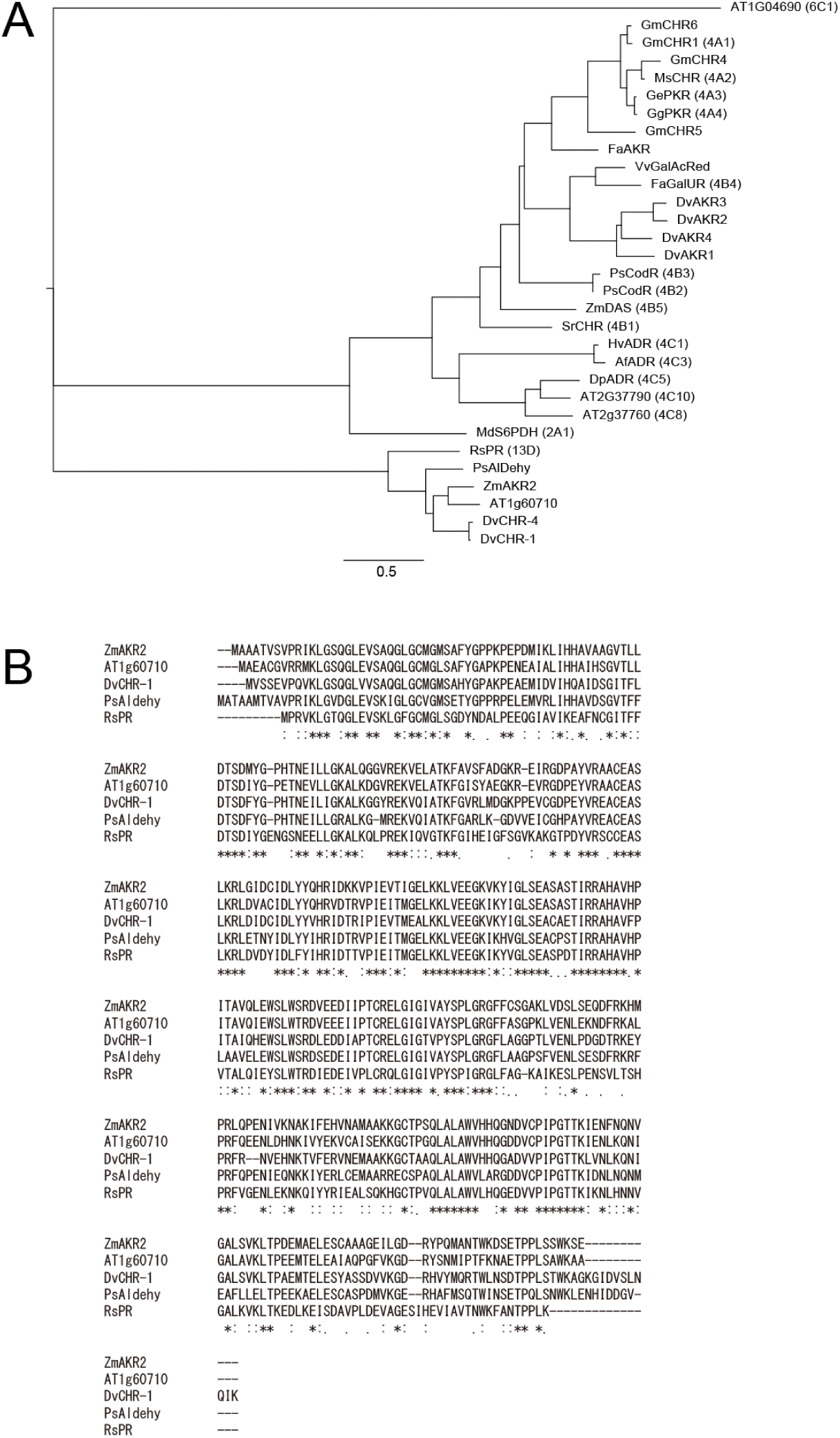
Phylogenetic analysis of DvCHR. A, Phylogenetic tree for plant Aldo-keto reductase superfamily protein. The entire amino acid sequences were aligned using ClustalW, and the tree was constructed by the Neighbor–Joining method. Bootstrap values of 1000 retrials are indicated on each branch, and the scale shows 0.1 amino acid substitutions per site. B, Comparison of putative amino acid sequence among *DvCHR* and its homologous genes.

Next, we analyzed the genomic sequence of *DvCHR*. *DvCHR* CDS was consisted of five exons and four introns. The genome structure was similar to *AT1G60710* which is also consisted of five exons and four introns but different from *GmCHR1* which is consisted of three exons and two introns (Fig. 5A). Since dahlia is considered as an octoploid, multiple alleles for *DvCHR* were expected. We identified four different *DvCHR* sequences in ‘SH’ and ‘RK’ based on the SNPs and indels (named *DvCHR-1* to *DvCHR-4*), one of four (*DvCHR-3*) has a deletion in the last exon and encoding truncated protein (Fig. 5B). We conducted inverse PCR to isolate *DvCHR* 5’ flanking sequence in ‘SH’ and ‘RK’, and we identified about 320 bp upstream from the start codon for *DvCHR-1* and *DvCHR-2*, and about 700 bp for *DvCHR-3* and *DvCHR-4*, however there was no sequence difference between ‘SH’ and ‘RK’.

**Fig. 5.**
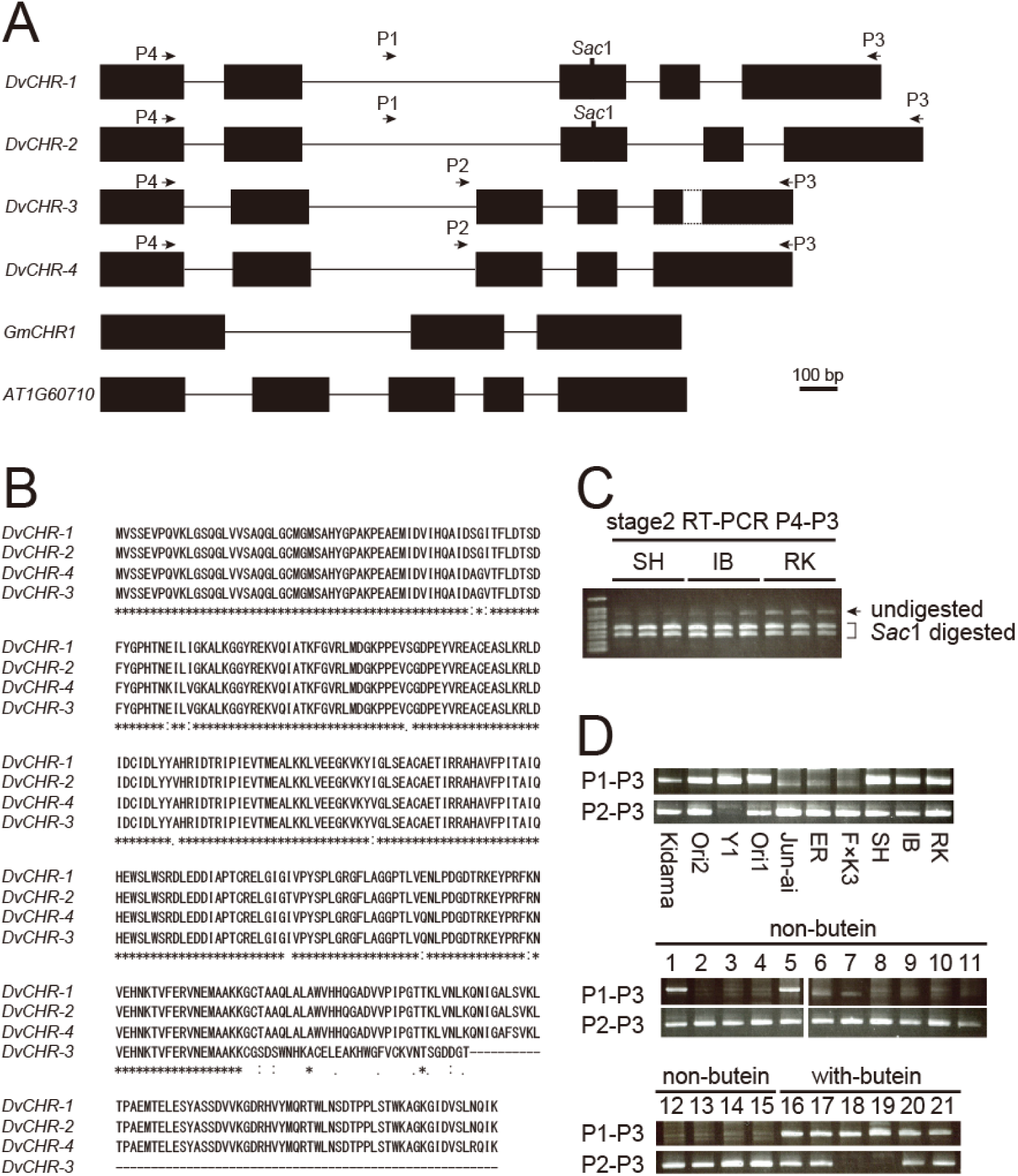
Analysis of *DvCHR* genetic background. A, Genomic structure of *DvCHR*. Black boxes indicate exons, and horizontal lines indicate introns. B, Alignment of putative amino acid sequences among DvCHR1-4. C, RT-PCR analysis of stage 2 ray florets using P4 and P3 primers and then digested with *Sac*I. *DvCHR-1* and *DvCHR-2* have *Sac*I digestion site while *DvCHR-3* and *DvCHR-4* do not. D, Genomic PCR analysis of *DvCHR* among butein accumulating cultivars and non-butein accumulating cultivars. 1: ‘Super Girl’, 2 ‘Yukino’, 3: ‘Cupid’, 4: ‘Atom’, 5: ‘Magokoro’, 6: ‘Saffron’, 7: ‘Gitt’s Attention’, 8: ‘Zannsetsu’, 9: ‘Hakuba’, 10: ‘Hakuyo’, 11: ‘Kokucho’, 12: ‘Fidalgo Blacky’, 13: ‘Ms. Noir’, 14: ‘Kazusa-shiranami’, 15: ‘Black Cat’, 16: ‘Yuino’, 17: ‘Matsuribayashi’, 18: ‘Red Velvet’, 19: ‘Michael J’, 20: ‘Suckle Pico’, 21: ‘Ittosei’.

To analyze which *DvCHR* gene is expressing in ‘SH’, ‘IB’ and ‘RK’, we performed RT-PCR combined with *Sac*I digestion using ray florets at stage 2. *DvCHR-1* and *DvCHR-2* have *Sac*I digestion site in the middle of the exon 3, while *DvCHR-3* and *DvCHR-4* do not. Though undigested band was a bit intense in ‘RK’ than ‘SH’, strong *Sac*I digested bands and faint undigested band were detected from all three cultivars, indicating *DvCHR-1* and *DvCHR-2* were much highly expressing than *DvCHR-3* and *DvCHR-4* (Fig. 5C).

Then, we analyzed the genetic background of *DvCHR* in other cultivars. We performed genomic PCR using P1 and P3 primers to detect *DvCHR-1* and *DvCHR-2* and using P2 and P3 primers to detect *DvCHR-3* and *DvCHR-4* (Fig. 5A). We, at first, used cultivars analyzed in Fig. 3C. Though both bands were detected from ‘Ori1’, P1-P3 bands were detected only from cultivars that accumulate butein and were not detected from cultivars that do not accumulate butein (Fig. 5D). To expand the experiment, we examined 21 more cultivars whose pigment composition was analyzed previously (Deguchi et al., 2013; Ohno et al., 2011a; 2011b; 2013b; 2021). Again, except for two cultivars (1: ‘Super Girl’ and 5: ‘Magokoro’), P1-P3 bands were detected from cultivars with butein but not from cultivars without butein (Fig. 5D). From these data, correlation between butein accumulation and genetic background of *DvCHR* was detected. Therefore, it was suggested that *DvCHR-1* and *DvCHR-2* are functional CHR in dahlia.

### Overexpression of either DvCHR-1, DvCHR-2, GmCHR5 or GmCHR6 is not sufficient for isoliquiritigenin accumulation in transgenic tobacco

To confirm the function of *DvCHR*, we produced *DvCHR-1* or *DvCHR-2* overexpressing transgenic plants in tobacco (*Nicotiana tabacum*) because *N. tabacum* is able to synthesize flavonoids (anthocyanins and flavonols) in flowers. We obtained several overexpression T_1_ lines, however, only flavonol was detected and isoliquiritigenin or butein were not detected in the flowers (Fig. 6C). We also produced *GmCHR5* or *GmCHR6* overexpression transgenic lines as a positive control, which are functional CHRs in soybeans (Mameda et al, 2018), however, we could not detect neither isoliquiritigenin nor butein in the flowers (Fig. 6C). These results suggested that single overexpression of *CHR* is not sufficient for isoliquiritigenin or butein accumulation in tobacco. We also produced *DvCHS2* or *DvCH3H* overexpression transgenic lines and crossed with *DvCHR-1*, *DvCHR-2* or *GmCHR5* overexpression lines to express multiple butein biosynthesis pathway genes. However, again only kaempferol was detected and neither isoliquiritigenin nor butein was detected in the flowers (Fig. 6C).

**Fig. 6.**
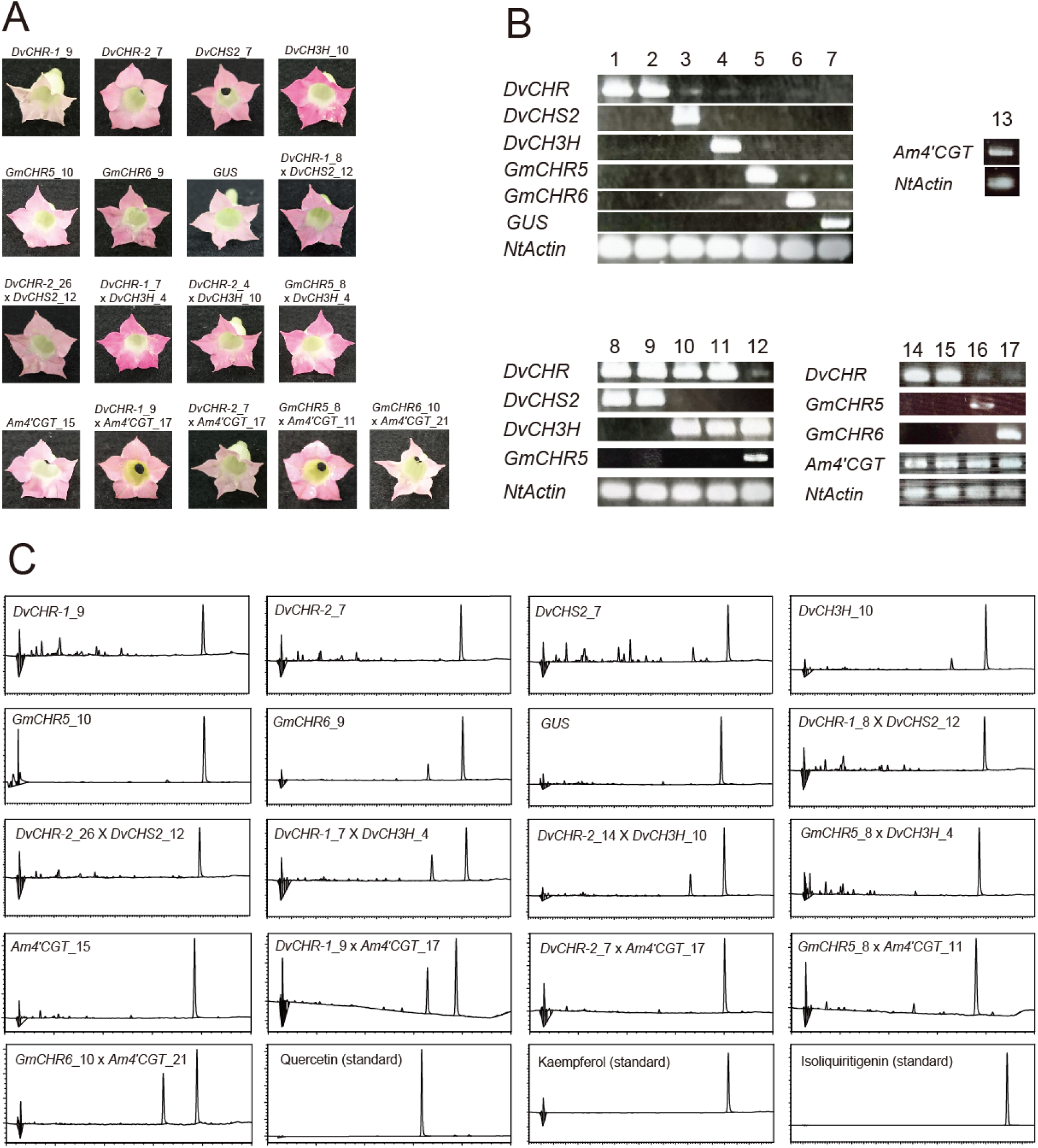
Analysis of overexpression transgenic *Nicotiana tabacum* plants. A, Photos of flowers of transgenic overexpression lines. B, Confirmation of transgene expression in flowers by RT-PCR. 1: *DvCHR-1*_9, 2: *DvCHR-2*_7, 3: *DvCHS2*_7, 4: *DvCH3H*_10, 5: *GmCHR5*_10, 6: *GmCHR6*_9, 7: *GUS*, 8: *DvCHR-1*_8 x *DvCHS2*_12, 9: *DvCHR-2*_26 x *DvCHS2*_12, 10: *DvCHR-1*_7 x *DvCH3H*_4, 11: *DvCHR-2*_14 x *DvCH3H*_10, 12: *GmCHR5*_8 x *DvCH3H*_4, 13: *Am4′CGT*_15, 14: *DvCHR-1*_9 x *Am4′CGT*_17, 15: *DvCHR-2*_7 x *Am4′CGT*_17, 16: *GmCHR5*_8 x *Am4′CGT*_11, 17: *GmCHR6*_10 x *Am4′CGT*_21. C, HPLC chromatogram analyzed at 380 nm in flowers.

### Transient co-overexpression of CHR, chalcone 4′-g lucosyltransferase and CaMYBA induced isoliquiritigenin accumulation in benthamiana tobacco leaves

Generally, flavonoids are accumulated as a glycoside *in vivo*, and dahlia accumulates isoliquiritigenin and butein as a 4′-glucoside or its derivatives (Harborne et al, 1990; Ohno et al., 2021). We assumed that tobacco plants are not able to accumulate isoliquiritigenin without glycosylation *in vivo* due to loss of suitable glucosyltransferase, thus, we tried to co-overexpress CHR (either *DvCHR-1*, *DvCHR-2*, *GmCHR5* or *GmCHR6*) with a snapdragon chalcone 4′-glucosyltransferase (*Am4′CGT*). We chose *Am4′CGT* because chalcone glucosyltransferase gene had not yet been isolated in dahlia, and the position of glycosylation by *Am4′CGT* was expected to be the same position for isoliquiritigenin 4′-glucoside or butein 4′-glucoside. To overexpress both CHR (either *DvCHR-1*, *DvCHR-2*, *GmCHR5* or *GmCHR6*) and *Am4′CGT*, we performed transient co-overexpression by agroinfiltration in *N. benthamiana* leaves. Again, we could not detect isoliquiritigenin in infiltrated leaves, however, we neither detect any other flavonoids (Fig. 7C), indicating flavonoid biosynthesis activity is very low in *N. benthamiana* leaves. To solve the problem, in addition to CHR (either *DvCHR-1*, *DvCHR-2*, *GmCHR5* or *GmCHR6*) and *Am4′CGT*, we co-overexpressed *CaMYBA*, a MYB transcription factor that positively regulates anthocyanin biosynthesis in pepper (Borovsky et al, 2004; Ohno et al., 2020) to intensify flavonoid biosynthesis activity in leaves. Eventually, isoliquiritigenin was detected from the *DvCHR* (*DvCHR-1* or *DvCHR-2*), *Am4′CGT* and *CaMYBA* co-infiltrated leaves (3/8 for *DvCHR-1* and 2/8 for *DvCHR-2*, Fig. 7C and D) and *GmCHR5* or *GmCHR6*, *Am4′CGT* and *CaMYBA* co-infiltrated leaves (3/8 for *GmCHR5* and 2/8 for *GmCHR6*, Fig. 7C and D), but not from *Am4′CGT* and *CaMYBA* co-infiltrated leaves (Fig. 7C). In addition, we detected delphinidin from the infiltrated leaves (Fig. S3). We also co-overexpressed *DvCHR-1* or *DvCHR-2* with *CaMYBA*, or *GmCHR5* or *GmCHR6* with *CaMYBA*, however, isoliquiritigenin was not detected (Fig. 7C). These results suggesting that *DvCHR* encodes chalcone reductase to synthesize isoliquiritigenin, and that chalcone glycosylation and another factor which is regulated by *CaMYBA* is required to accumulate isoliquiritigenin in tobacco leaves.

**Fig. 7.**
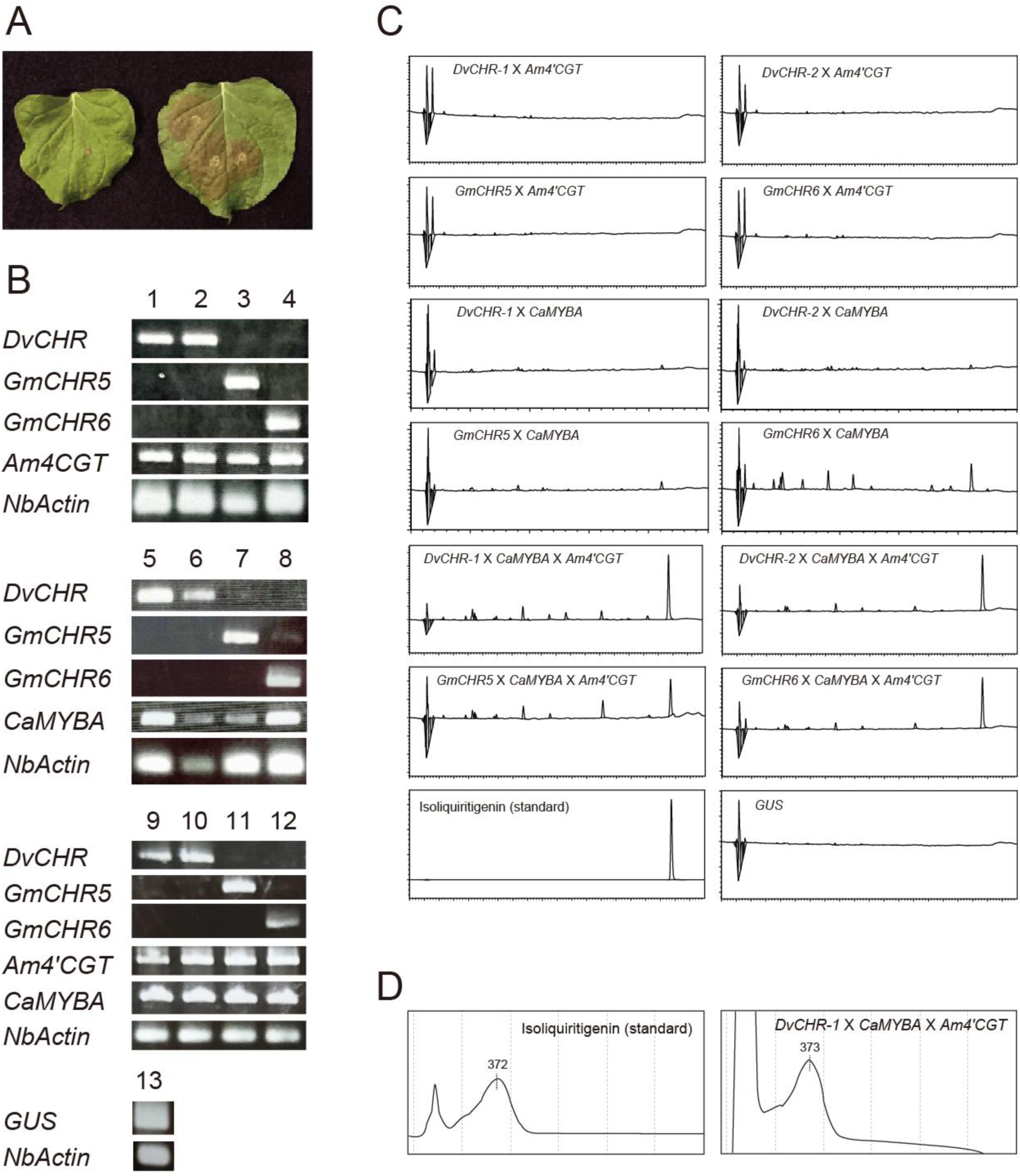
Analysis of transient overexpression analysis by agro-infiltration. A, Photos of infiltrated leaves. B, Confirmation of transgene expression in leaves by RT-PCR. 1: *DvCHR-1* x *Am4′CGT*, 2: *DvCHR-2* x *Am4′CGT*, 3: *GmCHR5* x *Am4′CGT*, 4: *GmCHR6* x *Am4′CGT*, 5: *DvCHR-1* x *CaMYBA*, 6: *DvCHR-2* x *CaMYBA*, 7: *GmCHR5* x *CaMYBA*, 8: *GmCHR6* x *CaMYBA*, 9: *DvCHR-1* x *CaMYBA* x *Am4′CGT*, 10: *DvCHR-2* x *CaMYBA* x *Am4′CGT*, 11: *GmCHR5* x *CaMYBA* x *Am4′CGT*, 12: *GmCHR6* x *CaMYBA* x *Am4′CGT*. C, HPLC chromatogram analyzed at 380 nm. D, Photodiode array analysis of detected peaks in isoliquiritigenin standard and *DvCHR-1* x *Am4′CGT* x *CaMYBA* triple overexpressing leaf.

We produced *Am4′CGT* overexpression transgenic tobacco lines and crossed with *DvCHR-1, DvCHR-2*, *GmCHR5* or *GmCHR6* overexpression lines to express both genes. However, isoliquiritigenin was not detected from flowers (Fig. 6C), indicating some gene(s) under *CaMYBA* regulation is essential for isoliquiritigenin accumulation in tobacco.

### DvCHR homologous gene expression was detected from other butein or aurone biosynthesizing Asteraceae species

Yellow flower color of *Asteraceae* species is divided into two types, one is derived from carotenoids and the other is derived from chalcones (butein and aurone). In snapdragon, aurone is directly synthesized from naringenin chalcone (Ono et al, 2006), however in *Asteraceae* species, aurone is synthesized through butein by polyphenol oxidase (Molitor et al, 2015; 2016). To analyze involvement of *DvCHR* homologous gene in chalcone accumulation in ray florets of other *Asteraceae* species, we performed hetero probe RNA gel blot analysis using a full length *DvCHR* cDNA. We chose dahlia, coreopsis, bidens and *Cosmos sulphureus* as a butein or aurone biosynthesis group, and chrysanthemum (*Chrysanthemum morifolium*), *Argyranthemum frutescens*, marigold (*Tagetes patula*), *Gaillardia* × *grandiflora* as a non-butein biosynthesis group. We detected intense band from a butein or aurone biosynthesis group but not from a non-butein biosynthesis group and snapdragon (Fig. 8). Therefore, clear correlation of *DvCHR* homologous gene expression and butein or aurone accumulation indicated *DvCHR* homologous gene expression is important for butein biosynthesis not only in dahlia but also in other butein accumulating *Asteraceae* species.

**Fig. 8.**
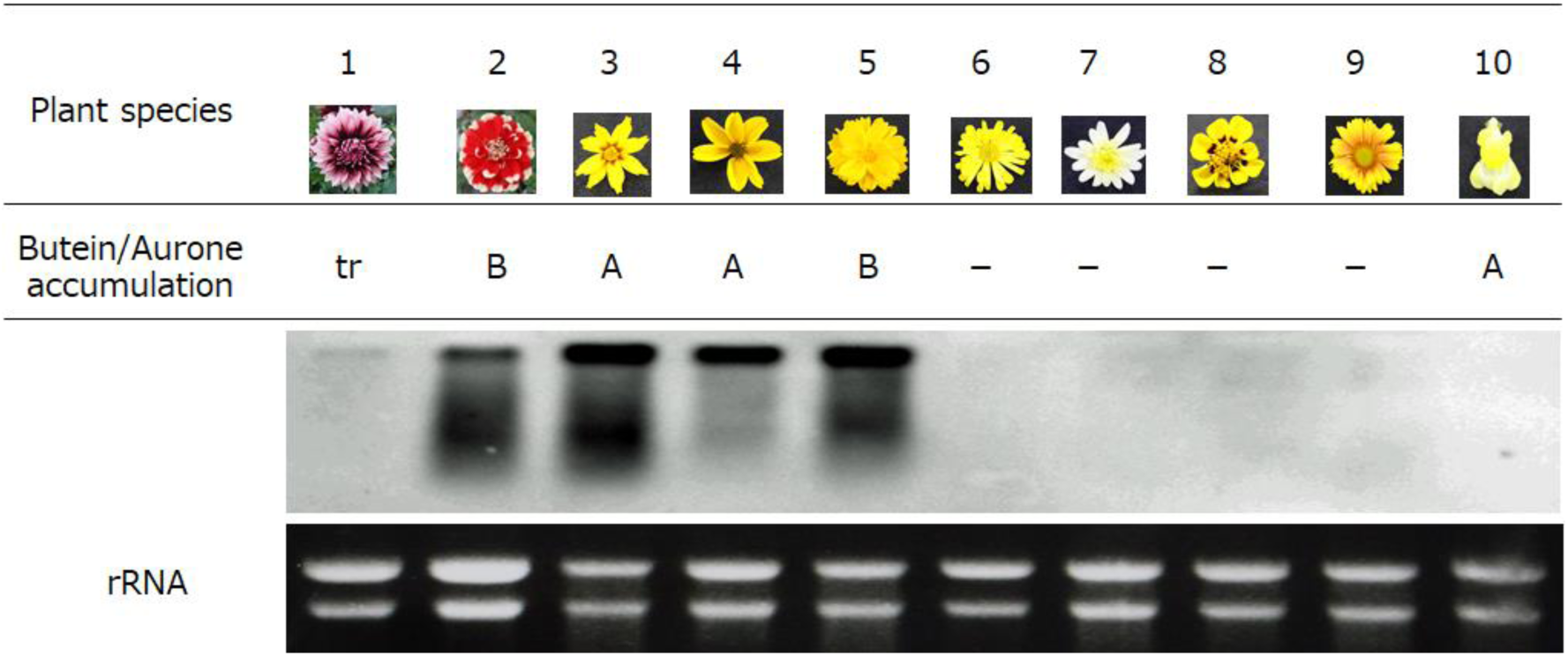
Hetero probe RNA gel blot analysis using *DvCHR* CDS as a probe. 1: *D. variabilis* ‘Kazusa-shiranami’, 2: *D. variabilis* ‘Shukuhai’, 3: *Coreopsis grandiflora* ‘Fairy Golden’, 4: *Bidens* ‘JuJu Gold’, 5: *Cosmos sulphureus*, 6: *Chrysanthemum morifolium* ‘Laub Fusha’, 7: *Argyranthemum frutescens* ‘Lemon Yellow’, 8: *Tagetes patula* ‘Harlequin’, 9: *Gaillardia* × *grandiflora* ‘Arizona Apricot’ 10: *Antirrhinum majus*. B indicates butein accumulation and A indicates aurone accumulation, and tr indicates trace amount of butein accumulation.

## Discussion

### DvCHR might be the determinant factor for isoliquiritigenin and butein biosynthesis

Butein is chemically described as 2′,3,4,4′-tetrahydroxychalcone, named after the genus *Butea*. Butein is found in limited species such as *Asteraceae* (dahlia, coreopsis), *Anacardiaceae* (*Searsia*) and *Fabaceae* (Semwal et al, 2015). As a flower color, butein derivative is an important compound because butein is one of a few flavonoids exhibiting bright yellow color. Many yellow flowers are exhibited yellow by carotenoids or betaxanthins, thus, plant species which are capable of biosynthesis only flavonoids rarely have yellow flower. For creation of a yellow flower by a transgenic approach in such species, to biosynthesize butein is much easier than to biosynthesize carotenoids or betaxanthins, thus, to reveal and to understand the butein biosynthesis pathway is quite important. The biosynthesis of butein is divided into two steps: the first step is isoliquiritigenin biosynthesis by CHS and CHR and the second step is conversion of isoliquiritigenin to butein by CH3H. In dahlia ray florets, butein is co-synthesized with isoliquiritigenin (Fig. 2; Ohno et al, 2011b; 2013a) suggesting the first isoliquiritigenin biosynthesis step is the determinant step. In a bicolor flower dahlia ‘Yuino’, post-transcriptional gene silencing of *DvCHS2* inhibits biosynthesis of not only anthocyanins and flavones but also isoliquiritigenin and butein (Ohno et al, 2011b, 2018a), suggesting *DvCHS2* is involved in isoliquiritigenin biosynthesis. On the other hand, ivory white cultivars where *DvCHS2* was expressed in ray florets, do not synthesize isoliquiritigenin and butein (Ohno et al, 2013b), suggesting *DvCHS2* is a requisite factor but not a sufficient factor. Thus, it was suggested that CHR is the determinant factor of isoliquiritigenin and butein biosynthesis in dahlia.

We isolated *DvCHR* from the comparative RNA-seq analysis and its expression showed correlation with butein accumulation among ‘SH’, ‘IB’, ‘RK’ and other cultivars (Fig. 3). Among four *DvCHR* sequences, all butein accumulating cultivars harbour *DvCHR-1* and/or *DvCHR-2*, while cultivars without *DvCHR-1* and/or *DvCHR-2* do not accumulate butein and isoliquiritigenin (Fig. 5D). These results indicate that *DvCHR* might be the determinant factor for isoliquiritigenin and butein biosynthesis. Lawrence (1931) conducted genetic analysis and proposed that the Y factor, which is dominant to y or I (ivory white) and behaves like tetrasomic, determines yellow color of octoploid dahlia. The Y factor has not been isolated for long time, but combined to our results, *DvCHR* would be the Y factor.

In ‘RK’, though expression level of *DvCHR* is lower compared to ‘SH’, but some extent of expression was still detected (Fig. 3B, Fig. 5C). We could not identify any mutation in *DvCHR* including promoter region, indicating *DvCHR* is unlikely to be causal mutation of flower color change. Expression level of *DvCHS2* was also reduced to about half by RNA-seq (Table S1), indicating loss of isoliquiritigenin and butein biosynthetic ability in ‘RK’ resulted that negative feedback regulation suppresses *DvCHR* and *DvCHS2* gene expression.

### DvCHR has been evolutionally differently developed from legume CHR

CHR that had been isolated so far is as an isoliquiritigenin biosynthesis enzyme in isoflavone biosynthesis pathway in legumes. In soybean, isoflavonoids are thought to play an important role as a phytoalexin in resistance to pathogens (Grahum et al, 2007, Sepiol et al, 2017), and as a *nod* gene inducer for nodule formation (Zuanazzi et al, 1998, Zhang et al, 2009). In the isoflavone biosynthetic pathway, CHR is required to synthesize 5-deoxy flavonoids (daizein, dihydroxyflavone). CHR is known as a multigene family in legumes, the soybean genome encodes 11 CHR paralogs (Mameda et al, 2018), among which GmCHR1, GmCHR5, GmCHR6 had been enzymatically characterized as functional CHR (Welle and Grisebach, 1988; Welle et al, 1991; Mameda et al, 2018). Legume CHRs belong to AKR 4A sub-superfamily (Jez et al, 1997; Bomati et al, 2005), and it was suggested CHR accepts coumaryl-trione as a substrate to synthesize isoliquiritigenin (Bomati et al, 2005). However, no other AKRs of other species belong to this 4A sub-family. In fact, we could not find any AKR 4A sub-family genes from RNA-seq of dahlia ray florets, suggesting CHR that belongs to AKR 4A sub-family is specific for legume isoflavonoid synthesis pathway. We identified four AKR 4B sub-family genes (*DvAKR1*-*DvAKR4*), but expression level of them did not correlate with butein accumulation (Fig. S2), indicating these genes are not involved in isoliquiritigenin biosynthesis in dahlia. Similar result was reported by Walliser et al (2021) that expression level of CHR-like genes (CHRl1-CHRl3) could not explain the butein biosynthesis.

*DvCHR*, which was isolated by comparative RNA-seq analysis between ‘SH’ and ‘RK’, also belongs to AKR family but phylogenetically far away from legume CHRs (Fig. 4A), which shares only 19% identity with GmCHR1 in putative amino acid sequence. DvCHR belongs to AKR13 family and phylogenetically locates near with *Arabidopsis* AT1G60710, *Zea mays* AKR2, *Rauvolfia serpentine* perakine reductase and *Perilla setoyensis* PsAKR. AKR superfamily enzymes are NAD(P)(H) binding oxidoreductases that metabolize a wide array of chemical substrates (Jez et al, 2001). PsAKR acts on the conversion of geraniol into citral and nerol, or perilla alcohol into perillaldehyde (Sato-Masumoto and Ito, 2010). *Rauvolfia serpentine* perakine reductase catalyzes an NADPH-dependent step in a side-blanch of the 10-step biosynthetic pathway of the alkaloid ajmaline (Sun et al, 2008). Here, perakine reductase displayed broad substrate acceptance including *p*-coumaryl aldehyde and cinnamic aldehyde (Sun et al, 2008). Thus, it may be possible that *DvCHR* accept *p*-coumaroyl CoA and malonyl-CoA as a substrate. *DvCHR* homologous gene expression was detected in petals of butein biosynthetic Asteraceae species, but not in non-butein biosynthetic Asteraceae species (Fig. 8), indicating *DvCHR* homologous genes function as a CHR in these species. These species belong to *Coreopsideae* tribe, indicating butein or aurone biosynthesis pathway by *DvCHR* homologous genes is shared in these species. Ecological importance of butein has not been revealed yet, however, phylogenetically close species are able to synthesize butein or aurone indicating these species has been developed CHR evolutionally different way from legume species.

### Co-overexpression of chalcone reductase, chalcone glucosyltransferase and CaMYBA is required for isoliquiritigenin accumulation in tobacco

Co-overexpression of either *DvCHR-1*, *DvCHR-2*, *GmCHR5* or *GmCHR6*, with *Am4′CGT* and *CaMYBA* successfully induced isoliquiritigenin accumulation in leaves, while co-overexpression of either *DvCHR-1*, *DvCHR-2*, *GmCHR5* or *GmCHR6*, with *CaMYBA* did not induce isoliquiritigenin accumulation (Fig. 7C). Am4′CGT was initially isolated as a UDP-glucose: chalcone 4′-O-glucosyltransferase for aurone formation (Ono et al, 2006), and silencing of *Am4′CGT* represses aurone production (Bradley et al, 2017). Enzymatic analysis revealed 2′, 4′, 6′, 4-tetrahydoroxychalcone was efficiently glycosylated and 2′, 4′, 6′, 3, 4-pentahydoroxychalcone was less efficiently glycosylated by Am4′CGT (Ono et al, 2006). Our results suggest that Am4′CGT is able to glycosylate 2′,4,4′-trihydroxychalcone (isoliquiritigenin) as well. In addition, our results indicate glycosylation is required to accumulate isoliquiritigenin *in vivo*. As far as we know, there is no study that reports transgenic tobacco successfully accumulated isoliquiritigenin by ectopic expression of legume CHRs. Instead, two studies reported successful accumulation of isoliquiritigenin in petunia flowers by introducing CHR. Davies et al (1998) reported that introduction of *MsCHR7* cDNA from *Medicago sativa* resulted in accumulation of butein 4-glucoside and butein 3-glucoside in flowers, isoliquiritigenin in pollen and trace amount of chalcones in leaves. Shimada et al (2006) reported that introduction of PKR1 from *Lotus japonicus* resulted in accumulation of trace amount of isoliquiritigenin in transformant flowers. Therefore, in contrast to tobacco, single overexpression of *CHR* is sufficient to accumulate isoliquiritigenin in petunia. This difference may be because endogenous genes such as glucosyltransferase enabled to accumulate butein or isoliquiritigenin in petunia, but tobacco does not have suitable set of genes.

Our results also indicating importance of ectopic expression of *CaMYBA* for isoliquiritigenin accumulation. We initially hypothesized that flavonoid biosynthesis activity is not enough in benthamiana leaves, because co-overexpression of *CHR* (either *DvCHR-1*, *DvCHR-2*, *GmCHR5* or *GmCHR6*) with *Am4′CGT* failed to accumulate flavonoid (Fig. 7C). However, transgenic tabacum plants expressing *CHR* (either *DvCHR-1*, *DvCHR-2*, *GmCHR5* or *GmCHR6*) with *Am4′CGT* also failed to accumulate isoliquiritigenin but accumulated kaempferol in flowers (Fig. 6C), indicating flavonoid biosynthesis activity is not the issue. Hence, another possibility is that the gene(s) promoted by CaMYBA is essential for isoliquiritigenin accumulation. CaMYBA belongs to AN2 clade MYB transcription factor and VIGS of *CaMYBA* indicated CaMYBA regulates *CaCHS*, *CaCHI*, *CaF3H*, *CaF3′5′H*, *CaDFR*, *CaANS*, *CaUFGT*, *CaANP* and *CaGST* (Zhang et al., 2015). Tobacco genome encodes floral tissue specific MYB transcription factor *NtAN2* (Pattanaik et al, 2010). Both CaMYBA and NtAN2 belongs to AN2 subgroup (Borovsky et al., 2004; Pattanaik et al, 2010), however, tobacco plants primally do not accumulate delphinidin suggesting set of genes regulated by CaMYBA and NtAN2 are different. Chen et al (2019) reported that grape hyacinth (*Muscari armeniacum*) *MaMYBA* overexpression tobacco plants accumulated delphinidin in leaves also indicating differential regulation of AN2 subgroup MYB transcription factors. In this study, it appears that ectopic expression of *CaMYBA* induced one or more gene expression and satisfied the condition to accumulate isoliquiritigenin. Further analysis is required to identify the gene that is essential for isoliquiritigenin accumulation in tobacco and future molecular breeding of yellow flowers by butein accumulation.

In conclusion, we identified a novel CHR that belongs to AKR 13 family and is phylogenetically far from known CHRs. In addition, we found that co-overexpression of *CHR*, *Am4′CGT* and *CaMYBA* is sufficient for isoliquiritigenin accumulation in tobacco. These results indicate essential role of *Am4′CGT* and *CaMYBA* in metabolic engineering of not only butein or aurone biosynthetic pathway but also of isoflavone biosynthetic pathway.

## Materials and methods

### Plant materials

The red-white bicolor dahlia cultivar ‘Shukuhai (SH)’ and its lateral mutant lines ‘Iwaibune (IB)’ and ‘Rinka (RK)’ were used for the experiment. ‘IB’ has dark red-white ray florets spontaneously occurred from ‘SH’, and ‘RK’ has purple ray florets spontaneously occurred from ‘IB’ (Fig. 2A). Ray florets were collected from field- or greenhouse-grown plants in the experimental field of Kyoto University (Kyoto, Japan) and used in the following analyses. In addition, seven cultivars or seedling lines (‘Evelyn Rumbold’, ‘Jun-ai’, ‘Kidama’, ‘FK3’, ‘OriW1’, ‘OriW2’ and ‘Y1’) grown in the same place were used for HPLC, gene expression and genotyping analyses.

### HPLC analysis

The pigment contents of ray florets were quantified with high-performance liquid chromatography (HPLC). Fresh ray florets were homogenized with a mortar and a pestle under liquid nitrogen, following which 1 mL of extraction solution (5 % hydrochloric acid in 50 % methanol) was added. For measurement of butein 4′-malonylglucoside, extraction solution (10 % acetic acid in 50 % methanol) was added instead of 5 % hydrochloric acid in 50 % methanol. The mixture was then centrifuged at 4°C at 15,000 rpm for 15 min, and the supernatant was collected and diluted 50 times with the same solvent. For hydrolysis, 1.2 mL of the diluted solution was boiled at 95°C for 2 h and 20 µL of the hydrolyzed solution was injected into the HPLC apparatus. The analysis was performed using an HPLC system (Hitachi L-7100, L-7200, L-7420, L-7500; Hitachi Systems, Ltd., Tokyo, Japan) or HPLC system (SCL-10AVP, SPD-M10AVP, CTO-10AVP. SIL-10ADVP, LC-10ADVP, FCV-10ALVP, DGU-14A, LCsolutions software; Shimazu Corp., Kyoto, Japan) with a C18 column (Nihon Waters K.K., Tokyo, Japan) that was maintained at 40°C. The detection wavelength was 350 or 380 nm for chalcones, and 530 nm for anthocyanidins. Eluant preparation and HPLC analysis proceeded according to Ohno et al. (2011b). Isoliquiritigenin (Tokyo Chemical Industry Co., Tokyo, Japan), butein (Tokyo Chemical Industry Co.), kaempferol (Wako Pure Chemical Industries), quercetin (Wako Pure Chemical Industries), pelargonidin (extracted by thin layer chromatography), cyanidin chloride (Wako Pure Chemical Industries, Osaka, Japan), delphinidin chloride (Nagara Science, Gifu, Japan), apigenin (Wako Pure Chemical Industries) and luteolin (Wako Pure Chemical Industries) were used as authentic standard. Quantification of flavonoids in ray florets was performed with three different ray florets.

### RNA-seq analysis

Total RNA of ‘SH’ and ‘RK’ were extracted from stage 2 ray florets (Fig. 2B) and two RNA samples each were mixed equally. Then the mixed RNA sample for each was sequenced by Illumina Hiseq2000 with 101-bp paired end, respectively. The number of total trimmed reads was 54,314,990 for ‘Shukuhai’ and 42,340,142 for ‘Rinka’. Each 4 data were *de novo* assembled by Trinity (Grabherr *et al*., 2011), and differentially expressed genes were analyzed by RSEM-based abundance estimation. Contigs of top 1000 FPKM values were chose to draw a scatter plot diagram.

### Isolation of genomic fragments and genotyping analysis

Genomic DNA was extracted from leaves or ray florets using MagExtractor Plant Genome (Toyobo, Osaka, Japan). Transcripts or genomic fragments were cloned to pTAC-1 vector (BioDynamics Laboratory Inc., Tokyo, Japan) for sequencing. Plasmids were extracted using Quick Gene Plasmid Kit S II (Kurabo, Osaka, Japan). For Inverse PCR, *Dra*I, *Eco*RV, *Hind*III, *Sac*I were used to obtain 5′ flanking sequence. Transcript sequences of *DvCHR*, *DvCHS2* and *DvCH3H* and genomic sequence of *DvCHR* were compared among ‘SH’, ‘IB’ and ‘RK’. Primers used for the analysis that were designed from RNA-seq data or known sequence data are shown in Table S2.

For genotyping of *DvCHR*, in addition to seven cultivars or seedling lines used for HPLC analysis, 21 cultivars (‘Super Girl’, ‘Yukino’, ‘Cupid’, ‘Atom’, ‘Magokoro’, ‘Saffron’, ‘Gitt’s Attention’, ‘Zannsetsu’, ‘Hakuba’, ‘Hakuyo’, ‘Kokucho’, ‘Fidalgo Blacky’, ‘Ms. Noir’, ‘Kazusa-shiranami’, ‘Black Cat’, ‘Yuino’, ‘Matsuribayashi’, ‘Red Velvet’, ‘Michael J’, ‘Suckle Pico’ and ‘Ittosei’) whose pigment composition was previously analyzed (Deguchi et al., 2013; Ohno et al., 2011a; 2011b; 2013b; 2021) were used. Genomic DNA was extracted with MagExtractor Plant Genome (Toyobo). PCR was performed with Blend Taq polymerase (Toyobo). The PCR was performed as follows: 94 °C for 2 min, followed by 30 cycles at 94 °C for 30 s, 55 °C for 30 s and 72 °C for 2 min. Primers used for the genotyping are shown in Table S3.

### Gene expression analysis

Total RNA was extracted from ray florets on different developmental stage using Sepasol RNA I Super G (Nacalai Tesque, Kyoto, Japan), purified with a high-salt solution for precipitation (Takara Bio Inc., Ohtsu, Japan) and reverse transcribed with ReverTra Ace (Toyobo). For qRT-PCR, ray florets were divided into 1-4 developmental stages (Fig. 2B; 1: early, 4: late), and relative expression level of *DvCHR* was investigated by qRT-PCR. Two µL of 50-fold diluted RT product was used as a template for qRT-PCR. qRT-PCR was performed with SYBR Premix Ex Taq™ II (Takara Bio Inc.) or THUNDERBIRD SYBR qPCR Mix (Toyobo) according to the manufacturer’s instructions using the LightCycler 480 system (Roche Diagnostics K.K., Tokyo, Japan). The qRT-PCR was performed as follows: 95°C for 5 min, followed by 45 cycles at 95°C for 10 s and 60°C for 30 s. Single-target product amplification was checked using a melting curve. The primers that were used for qRT-PCR are shown in Table S4.

For, RT-PCR analysis of *DvCHS2*, *DvCH3H*, *DvAKR1*, *DvAKR2*, *DvAKR3* and *DvAKR4*, total RNA was extracted from ray florets at stage2 using Sepasol RNA I Super G (Nacalai Tesque). After reverse transcription with ReverTra Ace (Toyobo), PCR was performed with Blend Taq polymerase (Toyobo). The PCR was performed as follows: 94 °C for 2 min, followed by 30-35 cycles at 94 °C for 30 s, 55 °C for 30 s and 72 °C for 2 min. Primers used for RT-PCR are shown in Table S5.

For analysis of transcript sequence of *DvCHR*, RT-PCR products of stage 2 ray florets using c25599_g2_i1 Fill-F and c25599_g2_i1 Fill-R primers (Table S2) were digested with *Sac*I (Takara Bio Inc). Reaction mixture was consisted of 2.5 µL of PCR product, 0.1 µL *Sac*I, 0.5 µL of 10 X L buffer and 1.9 µL water. The reaction mixture was incubated with 37 °C for overnight.

### Phylogenetic analysis

A phylogenetic tree was constructed based on the open reading frames or amino acids of various *AKR* genes and several other polyketide synthase genes using the Neighbor-Joining method (Saitou and Nei, 1987). The accession numbers for the amino acid sequences were as follows: 2A1: *Malus* x *domestica* NADP-dependent D-sorbitol-6-phosphate dehydrogenase (S6PDH) (P28475), 4A1: *Glycine max* CHR (CAA39261), 4A2: *Medicago sativa* CHR (CAA57782), 4A3: *Glycyrrhiza echinata* Polyketide reductase (PKR) (BAA12084), 4A4: *Glycyrrhiza glabra* PKR (BAA13113), 4B1: *Sesbania rostrata* CHR (CAA11226), 4B2: *Papaver somniferum* codeinone reductase (CodR) (AAF13739), 4B3: *Papaver somniferum* CodR (AAF13736), 4B4: *Fragaria* x *ananassa* D-galacturonate reductase (GalUR) (AAB97005), 4B5: *Zea mays* Deoxymugineic acid synthase (DAS) (BAF03164), 4C1: *Hordeum vulgare* Aldose reductase (ADR) (P23901), 4C3: *Avena fatua* ADR (Q43320), 4C5: *Digitalis purpurea* ADR (CAC32834), 4C8: *Arabidopsis thaliana* AT2g37760 (ABH07514), 4C10: *Arabidopsis thaliana* AT2G37790 (ABH07516), 6C1: *Arabidopsis thaliana* AT1G04690 (AAA87294), DvAKR1 (BDE26435), DvAKR2 (BDE26436), DvAKR3 (BDE26437), DvAKR4 (BDE26438), DvCHR-1 (BDE26431), DvCHR-4 (BDE26434), *Arabidopsis thaliana* AT1G60710 (OAP19317), *Fragaria x ananassa* AKR (AAV28174), *Glycine max* CHR4 (AIT97303), *Glycine max* CHR5 (NP_001353935), *Glycine max* CHR6 (BBC21043), *Perilla setoyensis* Alcohol Dehydrogenase (AlDehy) (AFV99150), *Rauvolfia serpentine* Perakine reductase (PR) (AAX11684), *Vitis vinifera* Galacturonic acid reductase (GalAcRed) (NP001268125) and *Zea mays* AKR2 (PWZ18047). The classification of each Aldo-keto reductase was according to Aldo-Keto Reductase (AKR) Superfamily homepage (https://www.med.upenn.edu/akr/).

### Production of stable overexpressing transgenic tobacco plants

*DvCHR-1*, *DvCHR-2*, *DvCHS2, DvCH3H, GmCHR5, GmCHR6* or *Am4′CGT* Cdna clone was at first subcloned to a pDONR221 (Invitrogen, Carlsbad, CA, USA) vector and then recombinated to a pGWB2 binary vector (Nakagawa et al, 2007) using Gateway system. *DvCHR-1*, *DvCHR-2*, and *DvCH3H* were cloned from ray florets of dahlia ‘Shukuhai (SH)’, *DvCHS2* was cloned from ray florets of dahlia ‘Yuino’ (Ohno et al, 2018a), *GmCHR5* and *GmCHR6* were cloned from bean sprouts of *Glycine max* and *Am4′CGT* was cloned from a yellow flower of *Antirrhinum majus*. Transgenic tobacco (*Nicotiana tabacum*) plants were obtained by standard *Agrobacterium tumefaciens* leaf disc transformation method according to Horsch et al (1988), using *Agrobacterium* strain EHA105. T_0_ transgenic plants which harbors one copy of the transgene were selected by genomic PCR, qPCR or DNA gel blot analysis. For genotyping, genomic DNA was extracted using SDS method. qPCR was performed as follows: 95°C for 2 min, followed by 40 cycles at 95°C for 10 s, 55°C for 5 s and 72°C for 20 s using THUNDERBIRD SYBR qPCR Mix (Toyobo). Single-target product amplification was checked using a melting curve. *NtActin* was used for an internal standard and copy number was calculated by relative quantification using standard curve. DNA gel blot analysis was conducted according to (Ohno et al, 2018a). T_0_ plants were self-crossed to obtain T_1_ generation, and T_1_ plants harboring transgene as homozygous were screened by genomic PCR and qPCR again. T_0_ plants or T_1_ plants were crossed to obtain multiple transgene overexpression lines. pGWB2-*GUS* overexpression lines were used as a mock treatment. Primers used for the gateway cloning are shown in Table S6, primers used for genotyping of transgenes are shown in Table S7 and primers used for transgene expression analysis are shown in Table S8.

### Agroinfiltration assay

To confirm gene function, transient expression of isoliquiritigenin and butein biosynthetic genes were examined in tobacco (*Nicotiana benthamiana*). cDNA of *DvCHR* (*DvCHR-1* and *DvCHR-2*), *DvCHS2, DvCH3H, GmCHR5, GmCHR6, Am4′CGT* and *CaMYBA* were subcloned to a pDONR221 (Invitrogen) vector and then recombinated to a pGWB2 binary vector (Nakagawa et al, 2007) using Gateway system. *CaMYBA* was cloned from purple flowers of *Capsicum annuum* ‘Peruvian Purple’ (Ohno et al, 2020). All constructs were transformed into the *A. tumefaciens* EHA105 strain. The *BETA-GLUCURONIDASE* (*GUS*) coding sequence was used for control assays.

Transient overexpression in benthamiana tobacco plants were performed according to Polturak et al (2016). Agrobacterium harboring transgene were infiltrated solely or co-infiltrated to 3-4 weeks seedling leaves. For co-infiltration, each agrobacteria suspension which has around 1.0 at OD_600_ were mixed in a 1:1 or 1:1:1 ratio before infiltration. Leaves used for subsequent pigment extraction and RT-PCR were sampled from 5-7 days post infiltration. Three biological replicates for each experiment were sampled, each consisting of at least three different leaves. The primers that were used for RT-PCR are shown in Table S8.

The *GUS* expression was confirmed by GUS staining. The *GUS* infiltrated leaves were infiltrated with GUS buffer (0.5 mM potassium ferricyanide, 0.5 mM potassium ferrocyanide, 0.3% TritonX-100, 5.0% methanol, 50 mM/pH 7.0 NaH_2_PO_4_ dissolved in water). Then, the leaves were infiltrated with GUS staining buffer (1.0 mM X-glucuronide, 0.5 mM potassium ferricyanide, 0.5 mM potassium ferrocyanide, 0.3% tritonX-100, 5.0% methanol, 50 mM/pH 7.0 NaH_2_PO_4_ dissolved in water) and leaved to stand at 37℃ overnight. Finally, the leaves were decolorized by infiltrating with 70% ethanol.

### Hetero probe RNA-gel blot analysis

Total RNA was extracted from flowers of *D. variabilis* ‘SH’ and ‘Kazusa-shiranami’, *Coreopsis grandiflora* ‘Fairy Golden’, *Bidens* ‘JuJu Gold’, *Cosmos sulphureus*, *Chrysanthemum morifolium* ‘Laub Fusha’, *Argyranthemum frutescens* ‘Lemon Yellow’, *Tagetes patula* ‘Harlequin’, *Gaillardia* × *grandiflora* ‘Arizona Apricot’ and *Antirrhinum majus* using Sepasol RNA I Super G (Nacalai Tesque) and purified with a high-salt solution for precipitation (Takara Bio Inc.). A five-micro gram of total RNA was used for the gel blot analysis. Full length CDS of *DvCHR* was labelled with the DIG RNA Labeling Kit (Sigma-Aldrich, St. Louis, USA). The probe was hybridized to the membrane at 50 °C overnight. Detection was conducted with CDP-Star (GE Healthcare Japan, Tokyo, Japan) and the chemiluminescence image was obtained using a LAS-3000 Mini (Fujifilm, Tokyo, Japan).

## Accession numbers

*DvCHR-1* cDNA (LC671883), *DvCHR-1* genomic sequence (LC671879), *DvCHR-2* cDNA (LC671884), *DvCHR-2* genomic sequence (LC671880), *DvCHR-3* cDNA (LC671885), *DvCHR-3* genomic sequence (LC671881), *DvCHR-4* cDNA (LC671886), *DvCHR-4* genomic sequence (LC671882), *DvAKR1* (LC671887), *DvAKR2* (LC671888), *DvAKR3* (LC671889), *DvAKR4* (LC671890) and *DvCH3H* (LC671891). Preliminary RNA-seq: ‘Shukuhai’ ray floret at stage 2 (DRR337625); ‘Rinka’ ray floret at stage 2 (DRR337626).

## Supplementary data

**Table S1.** FPKM values of flavonoid biosynthesis related genes between SH and RK stage 2 petals.

**Table S2.** Primers used for the isolation of *DvCHR*, *DvCHS2* and *DvCH3H*.

**Table S3.** Primers used for *DvCHR* genotyping.

**Table S4.** Primers used for real-time RT-PCR.

**Table S5.** Primers used for RT-PCR.

**Table S6.** Primers used for gateway cloning.

**Table S7.** Primers used for genotyping of transgenic tobacco plants.

**Table S8.** Primers used for varidation of transgene expression

**Fig. S1.** Comparison of putative amino acid sequence among *CH3H* and *F3′H* genes.

**Fig. S2.** RT-PCR analysis of *DvCHS2*, *DvCH3H*, *DvAKR1*, *DvAKR2*, *DvAKR3* and *DvAKR4* among SH, IB and RK stage2 petals.

**Fig. S3.** Detection of anthocyanins in *CHR*, *Am4′CGT* and *CaMYBA* co-infiltrated benthamiana leaves.

## Acknowledgements

This study was supported by the Grant-in-Aid for Young Scientists (No. 18K14455) from the JSPS to S.O.

## Author Contributions

S.O. conceived the study. S.O., M.H. and M.D. designed the experiments. S.O., H.Y., K.M., A.D., Y.K., M.Y., and F.T. conducted the experiments; S.O. drafted the manuscript. All authors approved the manuscript.

