## Supplemental figures and tables for "Identification of a novel *chalcone reductase* gene for isoliquiritigenin biosynthesis in dahlia (*Dahlia variabilis*)"

**Fig. S1**

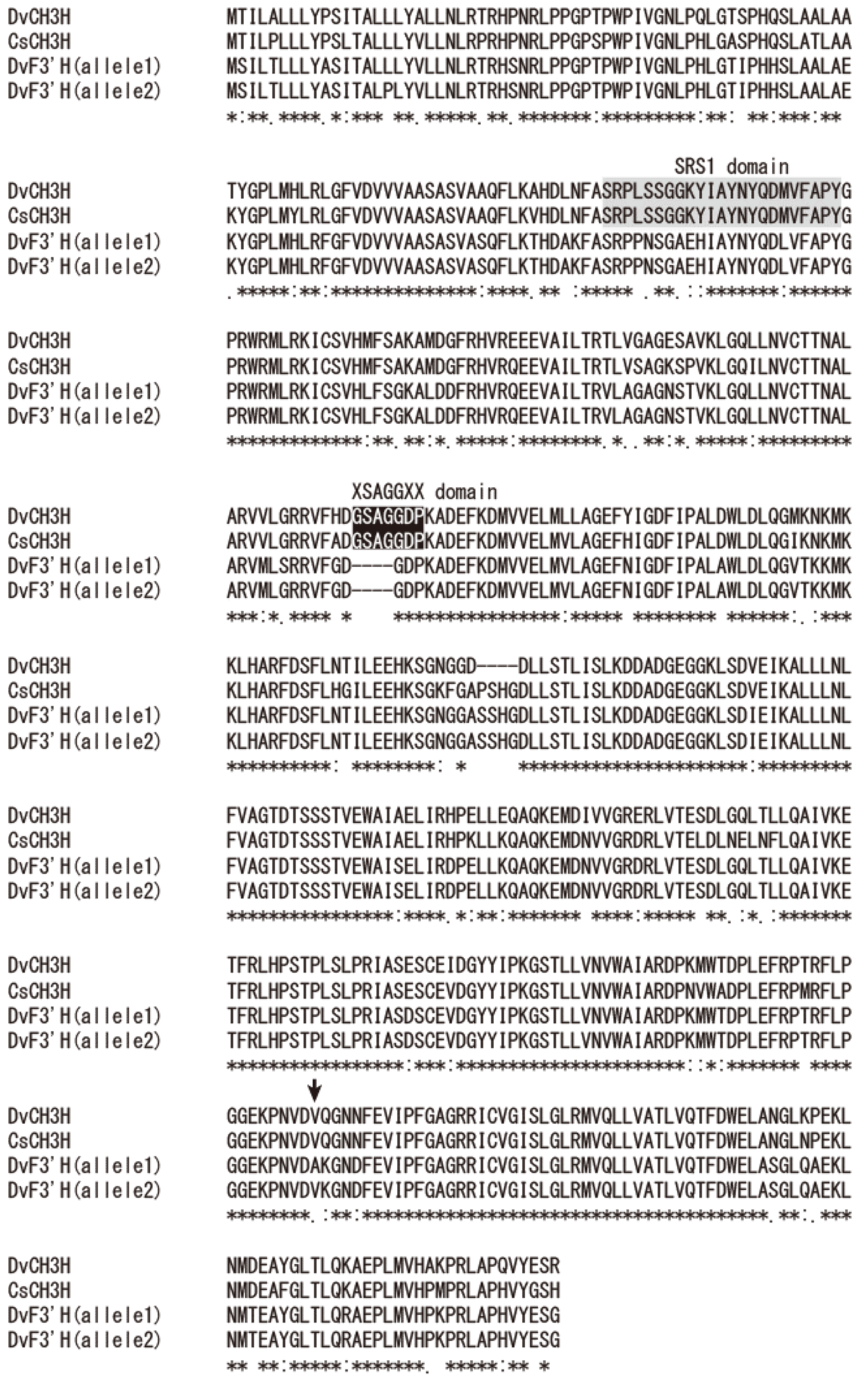


**Fig. S1.** Comparison of putative amino acid sequence among *CH3H* and *F3'H* genes. Gray box indicates SRS1 domain and black box indicates XSAGGXX domain, respectively. Allow indicates the valine residue at position 425 which is decisive for chalcone acceptance.

**Fig. S2**

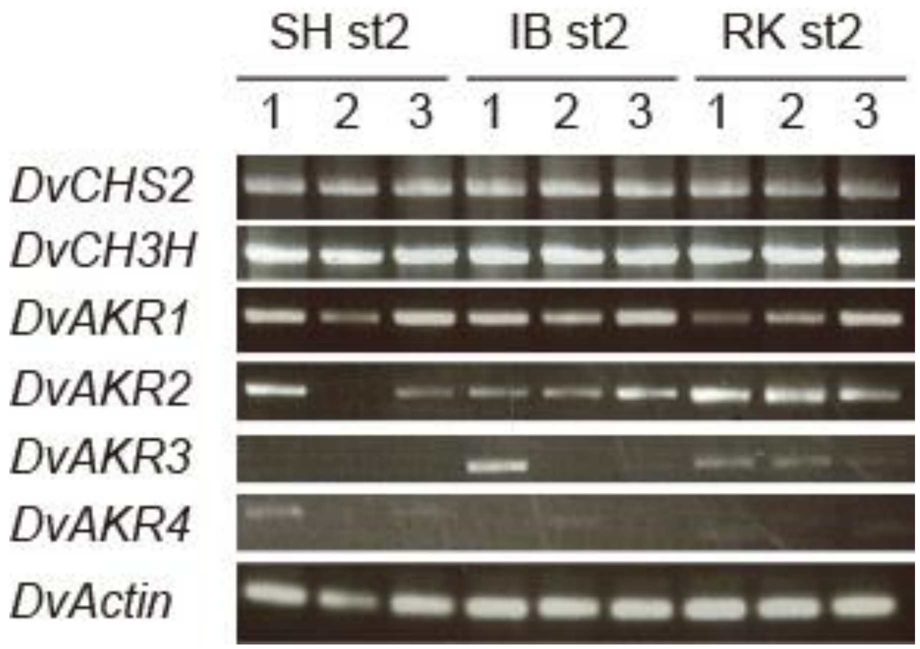


**Fig. S2.** RT-PCR analysis of *DvCHS2*, *DvCH3H*, *DvAKR1*, *DvAKR2*, *DvAKR3* and *DvAKR4* among SH, IB and RK stage2 petals.

**Fig. S3**

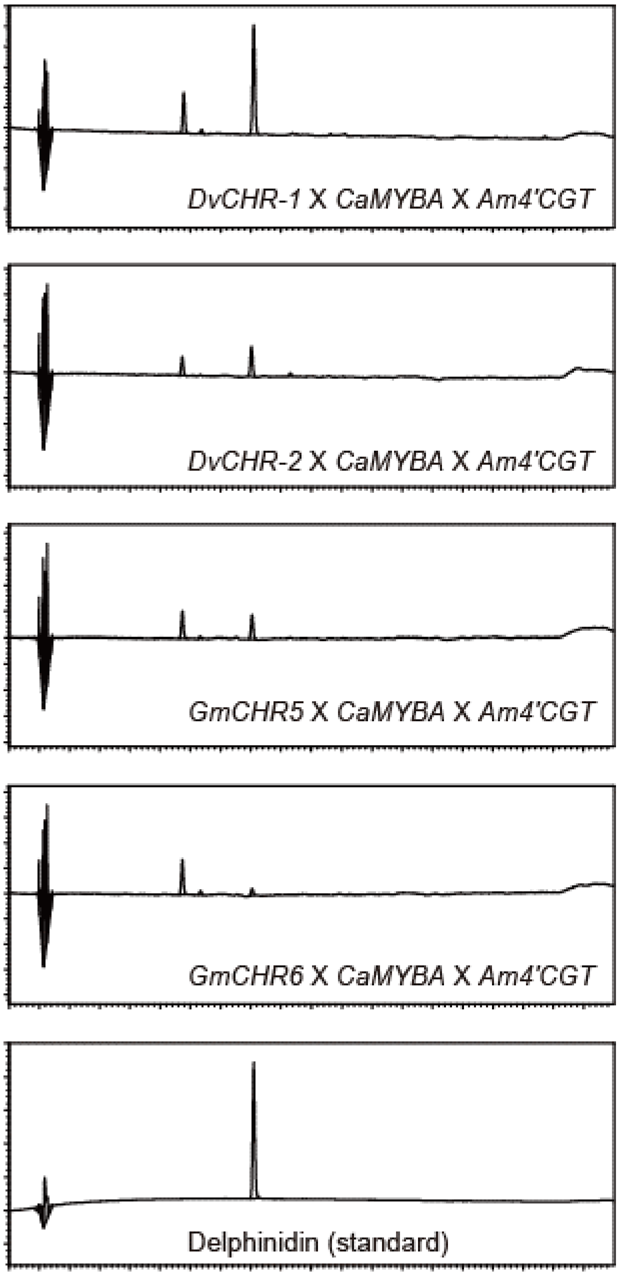


**Fig. S3.** Detection of anthocyanins in *CHR*, *Am4'CGT* and *CaMYBA* co-infiltrated benthamiana leaves. HPLC chromatogram analyzed at 530 nm.

Table S1. FPKM values of flavonoid biosynthesis related genes between SH and RK stage 2 petals.

| Protein | Name | SH FPKM | RK FPKM | SH/RK |
|---|---|---|---|---|
| CHS1 | c30947_g3_i1 | 487.34 | 296.92 | 1.64 |
| CHS2 | c32562_g1_i1 | 7502.34 | 3803.26 | 1.97 |
| CHI | c29501_g1_i1 | 1409.91 | 1329.54 | 1.06 |
| F3H | c34281_g1_i1 | 832.66 | 713.5 | 1.17 |
| DFR | c29663_g1_i1 | 1181.09 | 996.21 | 1.19 |
| ANS | c34758_g1_i1 | 1029.13 | 855.44 | 1.20 |
| 3GT | c37087_g1_i1 | 261.05 | 258.63 | 1.01 |
| 3MaT | c38195_g1_i1 | 171.63 | 175.64 | 0.98 |
| CH3H | c31660_g1_i2 | 139.9 | 137.26 | 1.02 |
| EFP | c27752_g1_i1 | 1127.68 | 1025.31 | 1.10 |
| FNS | c29165_g1_i1 | 595.29 | 512.29 | 1.16 |
| GT | c38674_g1_i1 | 477.38 | 401.15 | 1.19 |
| 3GRT | c37958_g1_i1 | 252.88 | 216.71 | 1.17 |
| MYB1 | c30720_g3_i1 | 860.43 | 687.39 | 1.25 |
| IVS | c33781_g1_i1 | 37.17 | 35.53 | 1.05 |
| CHR | c25599_g2_i1 | 2702.3 | 285.06 | 9.48 |

Table S2. Primers used for the isolation of *DvCHR*, *DvCHS2* and *DvCH3H*.

| Gene | Purpose | | Name | sequence (5'-3') |
|---|---|---|---|---|
| *DvCHR* | Sequence | | c25599_g2_i1 Full-F | CAACAGCAAAAACAACACACACTA |
| | Sequence | | c25599_g2_i1 Full-R | AAACACGCTCATAATCCACCCACA |
| | Sequence | | c25599_g2_i1-2R | CAAGATGGCAAAGAAACAACTTTTA |
| | Sequence | | c25599_g2_i1-3F | AGCTTGAAAAGACTCGATATCGACT |
| | Sequence | | c25599_g2_i1-SNPs1F | CCCACACCAACGAAATCCTCA |
| | Sequence | | c25599_g2_i1-SNPs2R | CGCCATCTCGTTCACTCTCTCAAAC |
| | Sequence | | c25599_g2_i1-SNPs3F | CCCACACCAACGAAATCCTCA |
| | Sequence | | c25599_g2_i1-type1F | AAAATTGATACGAGAGCCAGTGTGT |
| | Sequence | | c25599_g2_i1-type2F | TACGAGAGCCAGTGATTTTAACACC |
| | Sequence | | c25599_g2_i1-type34F | ATCAATTAATTAACAAGTGATTTCG |
| | Sequence | | c25599_g2_i1-6R | TCGTCCTCCAAATCTCGTGACCATA |
| | Sequence | | c25599_g2_i1_5'UTR-2F | TATGTAAAAAGGTTGAACATGGATT |
| | Sequence | | c25599_g2_i1_5'UTR-5F | ATTTTGAACTCTCCGCGCTAAGTAT |
| | Sequence | | c25599_g2_i1_Type1and2R | ACTAGGTCTTGGTCGATAGCTGATT |
| | Sequence | | c25599_g2_i1_Type2R | AGGTAATGCAAAAACACTGGTGTTA |
| | Sequence | | c25599_g2_i1_5'UTR-6F | CGCACTAAGTATACATGTTCATTTG |
| | Inverse PCR-F | EcoRV | c25599_g2_i1_Inv3F | TTCCTCGACACCTCCGACTTCTACG |
| | Inverse PCR-R | | c25599_g2_i1-Inv2R | AGTTTCACTTGTGGCACCTCGGATG |
| | Inverse PCR-F | DraI | c25599_g2_i1_Inv3F | TTCCTCGACACCTCCGACTTCTACG |
| | Inverse PCR-R | | c25599_g2_i1-Inv2R | AGTTTCACTTGTGGCACCTCGGATG |
| | Inverse PCR-F | SacI | c25599_g2_i1-Inv1F | AAGCACTGAAGAAACTGGTAGAGGA |
| | Inverse PCR-R | | c25599_g2_i1-4R | TCGCGATTTGAACCTTCTCTCTATA |
| | Inverse PCR-2nd-F | | c25599_g2_i1-Inv2F | AACTGGTAGAGGAAGGGAAAGTGAA |
| | Inverse PCR-2nd-R | | c25599_g2_i1-Inv2R | AGTTTCACTTGTGGCACCTCGGATG |
| | Inverse PCR-F | EcoRV | c25599_g2_i1-Inv5F | TTTGGAGTGCGATTGATGGACGGAA |
| | Inverse PCR-R | | c25599_g2_i1-Inv2R | AGTTTCACTTGTGGCACCTCGGATG |
| | Inverse PCR-F | EcoRV | c25599_g2_i1_Inv5F | TTTGGAGTGCGATTGATGGACGGAA |
| | Inverse PCR-R | | c25599_g2_i1-Inv3R | TATGTTTGAACATAGCAAGAGCTTG |
| | Inverse PCR-F | DraI | c25599_g2_i1_Inv4F | ACACCAACGAAATCCTCATCG |
| | Inverse PCR-R | | c25599_g2_i1-Inv3R | TATGTTTGAACATAGCAAGAGCTTG |
| | Inverse PCR-F | EcoRV | c25599_g2_i1_Inv4F | ACACCAACGAAATCCTCATCG |
| | Inverse PCR-R | | c25599_g2_i1_Inv4R | TAAATATTGAGTACTCACTATCCTA |
| | Inverse PCR-F | HindIII | c25599_g2_i1_Inv8F | CTAAGACCTACTCGCAAAACGGTCG |
| | Inverse PCR-R | | c25599_g2_i1-Inv1R | ACCTTGTGAACCTAGTTTCACTTGT |
| | Inverse PCR-2nd-F | | c25599_g2_i1_Inv9F | AACGGTCGCAACTTTGTGACCGCAT |
| | Inverse PCR-2nd-R | | c25599_g2_i1-Inv2R | AGTTTCACTTGTGGCACCTCGGATG |
| | Inverse PCR-F | HindIII | c25599_g2_i1_Inv9F | AACGGTCGCAACTTTGTGACCGCAT |
| | Inverse PCR-R | | c25599_g2_i1-Inv2R | AGTTTCACTTGTGGCACCTCGGATG |
| *DvCHS2* | RT-PCR | | CHS2-Full F | TCTTATTACTGCTCGCAATATCTT |
| | RT-PCR | | CHS2-Full R | AGTTAGGGCGAAATCGGCATGGTA |
| *DvCH3H* | 3'Race | | CH3H-3'RACE | CTCTCCCTACCAAGAATTGCATC |
| | 3'Race | | CH3H-3'RACE-Nested | GGACGGACCCACTTGAATTC |
| | 5'Race | | CH3H-5'RACE | GCTAATACCCACACAAATCCTTC |
| | 5'Race | | CH3H-5'RACE-Nested | GGAATTCAAGTGGGTCCGTC |
| | RT-PCR | | CH3H Full-F | GAAAATTAAACCACCCACAC |
| | RT-PCR | | CH3H Full-R | CTCGAGTTTATTAATTATGC |
| | Sequencing | | CH3H 121F | GTCGGAAACCTACCGCAAC |
| | Sequencing | | CH3H 144F | CACCAGTCCGCACCAATCG |
| | Sequencing | | CH3H 381F | GAAGATTTGCTCGGTGCATATG |
| | Sequencing | | CH3H 505F | GGTCAACTGCTTAACGTG |
| | Sequencing | | CH3H 920F | CAGTGGAATGGGCAATAGCCG |
| | Sequencing | | CH3H 1514R | TAGACTTGAGGAGCTAGCCTCG |

Table S3. Primers used for *DvCHR* genotyping.

| Genes | Forward primers | Reverse primers |
|---|---|---|
| *DvCHR-1,-2* | ACAACGATTTGCGACAGAAATCATA | GTCCTTTGCATATACACATGCCTAT |
| *DvCHR-3,-4* | ATCAATTAATTAACAAGTGATTTCG | GTCCTTTGCATATACACATGCCTAT |

Table S4. Primers used for real-time RT-PCR.

| Genes | Forward primers | Reverse primers |
|---|---|---|
| *DvCHR* | ACAAGGAAGGAATATCCGAGGTTTA | TGGTGCACCCACGCTAACGCTAGTT |
| *DvActin* | TAAGAGCGGCAGACATTGGGATATT | ACTTCTTCACAACCACTCTCCACTA |

Table S5. Primers used for RT-PCR.

| Genes | Forward primers | Reverse primers |
|---|---|---|
| *DvCHS2* | TCTTATTACTGCTCGCAATATCTT | AGTTAGGGCGAAATCGGCATGGTA |
| *DvCH3H* | ATGACTATTCTAGCCCTACTACTCT | TTAACGACTTTCATAGACTTGAGGA |
| *DvAKR1* | CAACCCAAGAAAATAAGGTTAGAG | GTGTTACATCTTTGGTGCAAAATCA |
| *DvAKR2* | GGCAACCATTCCGGACAC | AGTGCTCAAAGTTACTCTCTTGA |
| *DvAKR3* | AAACCACTAACAAAGTTCCC | CACAATATTCAATCCACAAAACAC |
| *DvAKR4* | TATTCATTCCAGAGACAACGA | GCTACGATACATCCAAACCGA |
| *DvActin* | TAAGAGCGGCAGACATTGGGATATT | ACTTCTTCACAACCACTCTCCACTA |

Table S6. Primers used for gateway cloning.

| Gene | Name | sequence (5'-3') |
|---|---|---|
| *DvCHR* | c25599_g2_i1_gateway-F2 | AAAAAGCAGGCTCTATGGTCTCATCCGAGGTGCCACAAG |
| | c25599_g2_i1_gateway-R2 | AGAAAGCTGGGTCTCACTTAATTTGGTTTAACGAAACA |
| *DvCH3H* | CH3H-gateway-F2 | AAAAAGCAGGCTCTATGACTATTCTAGCCCTACTACTCT |
| | CH3H-gateway-R2 | AGAAAGCTGGGTCTTAACGACTTTCATAGACTTGAGGA |
| *DvCHS2* | DvCHS2_gateway-F | AAAAAGCAGGCTCTATGGCATCTTCGGTCGATATTGCCG |
| | DvCHS2_gateway-R | AGAAAGCTGGGTCTTAGGGCGAAATCGGCATGGTAGTT |
| *GmCHR5* | Gateway_GmCHR5-Full_F | AAAAAGCAGGCTCTATGGCTGCCACCACCTTAGT |
| | Gateway_GmCHR5-Full_R | AGAAAGCTGGGTCTCATTCTTCATCCCATAGAT |
| *GmCHR6* | Gateway_GmCHR6-Full_F | AAAAAGCAGGCTCTATGGCGGCTGCTATTGAAAT |
| | Gateway_GmCHR6-Full_R | AGAAAGCTGGGTCTTATATTTCATCATCCCAGA |
| *Am4'CGT* | Gateway_Am4'CGT-Full_F | AAAAAGCAGGCTCTATGGGAGAAGAATACAAGAA |
| | Gateway_Am4'CGT-Full_R | AGAAAGCTGGGTCTTAACGAGTGACCGAGTTGA |
| *CaMYBA* | CaMYBA-attB-F | AAAAAGCAGGCTTCATGAATACTGCTATTATTGC |
| | CaMYBA-attB-R | AGAAAGCTGGGTACTAATTAAGTAGATTCCATA |
| | attB1 adapter primer | GGGGACAAGTTTGTACAAAAAAGCAGGCT |
| | attB2 adapter primer | GGGGACCACTTTGTACAAGAAAGCTGGGT |
| | pGWB2_insertcheck-F | CTCCACTGACGTAAGGGATGAC |
| | pGWB2_insertcheck-R | TTCGAGCTCTAAGCGCTGTT |
| | NPTII_F | ATGGGGATTGAACAAGATGGA |
| *NPTII* | NPTII_396F | CCCATTCGACCACCAAGCGAAACAT |
| | NPTII_R | TCAGAAGAACTCGTCAAGAAG |

Table S7. Primers used for genotyping of transgenic tobacco plants

| Gene | Purpose | sequence (5'-3') | Reference |
|---|---|---|---|
| *DvCHR* | qPCR | ACAAGGAAGGAATATCCGAGGTTTA | |
| | qPCR | TGGTGCACCCACGCTAACGCTAGTT | |
| *DvCH3H* | qPCR | GCAGTTTTTAAAAGCTCATGATCT | |
| | qPCR | GGTGCGAACACCATATCTTGATAA | |
| *DvCHS2* | qPCR | AAGGTTTGGATTGGGGTGTTCTGTT | |
| | qPCR | TTAGGGCGAAATCGGCATGGTAGT | |
| *GmCHR5* | qPCR | AATTGCATCCAGAGATCAAC | Mameda et al. (2018) |
| | qPCR | TTTCCAGGCTTAGTAGCAATG | |
| *GmCHR6* | qPCR | GGAGAATGATGTGCTGAAAG | |
| | qPCR | AACGGCTCTGATATATTTCATC | |
| *Am4'CGT* | qPCR | GGACATAGCGTTGGCTACCCGTGAT (To amplify NPTII) | |
| | qPCR | TCAGAAGAACTCGTCAAGAAG (To amplify NPTII) | |
| | Genomic PCR | AAAAAGCAGGCTCTATGGGAGAAGAATACAAGAA | |
| | Genomic PCR | AGAAAGCTGGGTCTTAACGAGTGACCGAGTTGA | |
| *NtActin* | qPCR | GGACATAGCGTTGGCTACCCGTGAT | |
| | qPCR | TCAGAAGAACTCGTCAAGAAG | |

Table S8. Primers used for varidation of transgene expression.

| Gene | Purpose | sequence (5'-3') | Reference |
|---|---|---|---|
| *DvCHR* | RT-PCR | ATGGTCTCATCCGAGGTGCCACAAG | |
| | RT-PCR | TCACTTAATTTGGTTTAACGAAACA | |
| *DvCH3H* | RT-PCR | ATGACTATTCTAGCCCTACTACTCT | |
| | RT-PCR | TTAACGACTTTCATAGACTTGAGGA | |
| *DvCHS2* | RT-PCR | ATGGCATCTTCGGTCGATATTGCC | |
| | RT-PCR | TTAGGGCGAAATCGGCATGGTAGT | |
| *GmCHR5* | RT-PCR | AATTGCATCCAGAGATCAAC | Mameda et al. (2018) |
| | RT-PCR | TTTCCAGGCTTAGTAGCAATG | |
| *GmCHR6* | RT-PCR | GGAGAATGATGTGCTGAAAG | |
| | RT-PCR | AACGGCTCTGATATATTTCATC | |
| *Am4'CGT* | RT-PCR | TGGGAGAAGAATACAAGAAAACA | Hoshino et al. (2019) |
| | RT-PCR | ACGAGTGACCGAGTTGATGA | |
| *CaMYBA* | RT-PCR | AAAAAGCAGGCTTCATGAATACTGCTATTATTGC | |
| | RT-PCR | AGAAAGCTGGGTACTAATTAAGTAGATTCCATA | |
| *GUS* | RT-PCR | ATGGTCCGTCCTGTAGAAACCCCAA | |
| | RT-PCR | TTATTGTTTGCCTCCCTGCTGCGGT | |
| *Actin* | RT-PCR | GGACATAGCGTTGGCTACCCGTGAT | |
| | RT-PCR | TCAGAAGAACTCGTCAAGAAG | |
